# The scents of summer: Plant-diversity and vegetation-height drive health-relevant scent exposure in urban greenspaces

**DOI:** 10.64898/2026.09.22.753177

**Authors:** William T. Kay, Kieran E. Storer, Molly Tucker, Katherine Willis

## Abstract

Urban green infrastructure is increasingly promoted as a public-health intervention, yet the biological features that determine exposure to potentially beneficial plant-derived volatile compounds remain poorly understood. Here we quantified airborne terpenes and terpenoids across five contrasting urban greenspaces in Oxford, UK, and related chemical profiles to plant diversity, vegetation structure and surrounding habitat characteristics. Ambient air samples collected across spring and summer were analysed by thermal-desorption gas chromatography–mass spectrometry, alongside field surveys of taxonomic richness, canopy cover, vegetation volume and ground composition. Terpene and terpenoid scentscapes differed strongly among sites and through time, with monoterpenes dominating detected profiles. Sites with greater plant richness and greater vegetation volume below 1.5 m consistently showed higher terpene/terpenoid abundance, whereas total vegetation volume, canopy cover and tree abundance were weaker and less consistent predictors. These relationships were specific to plant-associated chemical classes and were not observed for predominantly anthropogenic volatile compounds. These patterns were resolved across only five independent sampling sites, and the relationships reported here should therefore be considered exploratory. Our findings suggest that human exposure to plant-derived volatiles in cities may be maximised not simply by increasing tree cover, but by designing biodiverse, structurally complex lower-to-mid-storey vegetation within the breathing zone. These results provide an empirical basis for incorporating scentscape ecology into health-oriented urban greenspace design.

## Introduction

The evidence linking natural environments to positive effects on human health is substantial and well established. ^1–3^. A large body of evidence has been provided both at the psychological level, with benefits such as the reduction of stress, anxiety and fatigue ^4–6^ and at the physiological level, with positive immunomodulatory, analgesic, neuroprotective, anti-inflammatory, respiratory, anticancer, and antimicrobial effects cited ^7–11^. These benefits have been linked to all the human senses with visual features (such as the level of greenness), sounds (such as birdsong), and scents (such as terpenes) having all been shown to individually improve human health ^9,12–18^.

Such is the healing power of nature that urban green infrastructure is becoming increasingly promoted as a solution to health-related challenges in cities. For example, in the UK, the 2023 Green Infrastructure Framework from Natural England aims to support the creation of “good quality green infrastructure for people and for nature” ^19^. Similarly, the WHO have published ‘Urban green spaces and health’ which concludes the need for both small and large biologically diverse green spaces in urban areas ^20^. Indeed, many urban greenspaces have already been designed with the aim of achieving benefits such as these – a relatively modern area of study termed biophilic design ^18,21,22^.

But which urban greenspaces are most effective in promoting and restoring health? and which properties of these greenspaces are most important for maximising health gains? and how can answering this help us to design and build spaces more beneficial to health?

In terms of scent, accounting for the scentscape (the total profile of volatile compounds in a specific time and place) for urban greenspace design purposes has been widely discussed as a concept ^23–28^. Plants constitutively release biogenic volatile organic compounds (bVOCs), and consequently urban areas can be rich in these compounds ^29^. In particular, it is the terpenes/terpenoids which have been found to impart most benefit through lowering cortisol, decreasing blood pressure and heart rate, and improving mood, cognition, and immune function ^2,30–32^. Taufer et al ^11^ reviewed primary papers linking human health to urban greenspaces scents and noted that “remarkably” every paper found some positive effect.

Recent research has shown that, in terms of terpene/terpenoid presence, not all urban greenspaces are equal ^8,33,34^, yet to date, empirical work on community-level bVOC profiles and how these relate to the characteristics of a space and the biodiversity within it have remained largely theoretical ^35^. Terpenoid emissions are well-known to be induced by herbivory and pathogen attack ^36,37^, and are modified by abiotic stresses including drought, heat and UV ^38,39^. The diversity of plants, in terms of the numbers, vegetation volume, and diversity, will also be major factors in terpene/terpenoid presence ^40–44^. Indeed, this has been shown in an experimental grassland system where increases in plant species richness positively correlated with terpene/terpenoid detection ^35^. Vegetation structure and height may further affect the spatial and temporal availability of natural compounds such as terpenes ^44^. For example, terpenes from urban trees may be dispersed upwards and away from the human nose, whereas the compounds from lower vegetation may be more available for inhalation ^45^.

To assess which natural characteristics are most important for terpene/terpenoid presence and prevalence, this study asks three questions: firstly, which terpenes and terpenoids are present among five contrasting urban greenspaces in Oxford, UK; secondly, whether airborne terpene and terpenoid profiles differ between sites; and finally, which physical and vegetative features best explain these differences. We hypothesise that sites with greater plant diversity, more tree cover, more biomass, and more structurally natural habitat will show higher terpene/terpenoid presence and prevalence.

## Methods

This study used a comparative field-based design across five contrasting urban greenspaces in and around Oxford, UK, to investigate how site characteristics relate to airborne terpene and terpenoid presence. At each site, we first characterised the vegetation and habitat structure by surveying plant diversity and estimating features such as tree cover and ground composition. We then collected ambient air samples for chemical analysis using thermal-desorption-gas-chromatography-mass-spectrometry (TD-GC-MS) to identify terpene and terpenoid compounds. Finally, we used linear modelling to test whether variation in terpene/terpenoid presence and prevalence could be explained by differences in biodiversity and habitat structure across sites.

### Site locations and surveying

Five outdoor urban greenspace sites around Oxford with different ecologies (Figure 1) were used as sampling sites: Oxford Botanic Garden, University Parks (Dense tree area), University Parks (Open grassy area), Warneford Meadow, and Wytham Woods ^84^. For all outdoor spaces, above-ground family, genus, and species counts were calculated using quadrats. A 20 m diameter circle was measured from the bVOC sampling site and ten random coordinates were chosen for 1m x 1m quadrats. Species were identified using the Seek app, and cross-referenced with a physical key ^85^.

**Figure 1:**
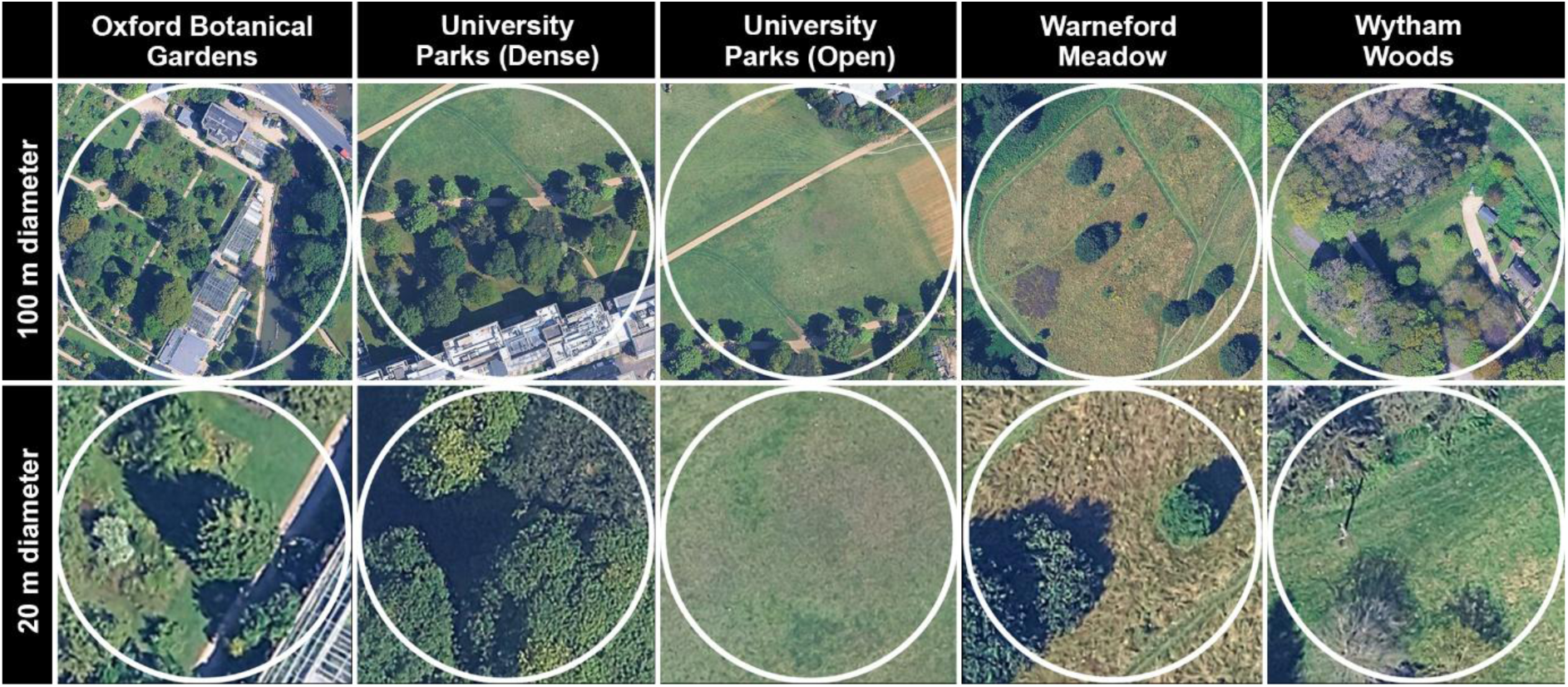
Aerial imagery of the five outdoor sampling sites used in this study: Oxford Botanic Garden (BO), University Parks Dense (UD), University Parks Open (UO), Warneford Meadow (WA), and Wytham Woods (WY). For each site, the upper panel shows a 100 m diameter, while the lower panel shows a 20 m diameter (white circles). Map data: Google Earth Pro, Infoterra Ltd & COWI / Maxar Technologies (Imagery date: April 30, 2024).

Vegetation volume was estimated using multiple methods. For grasses and ground level cover, vegetation height was measured in 10 random points, averaged, and multiplied by the area of grass coverage within the 20 m diameter. For flowerbeds and other non-grass mid height vegetation, Polycam (poly.cam) 3D scanning and spatial capture app was used on a Google Pixel 9A (example imagery in Figure 2). After cropping and resizing, vegetation volume below 1.5 m was calculated using the volume tool. For trees, canopy widths were measured using a laser measure tool - height was extrapolated from photos using Adobe Photoshop ® and envelope volume was using an ellipsoidal approximation: 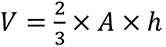, where *A* is the projected canopy area and ℎ is canopy height.

**Figure 2:**
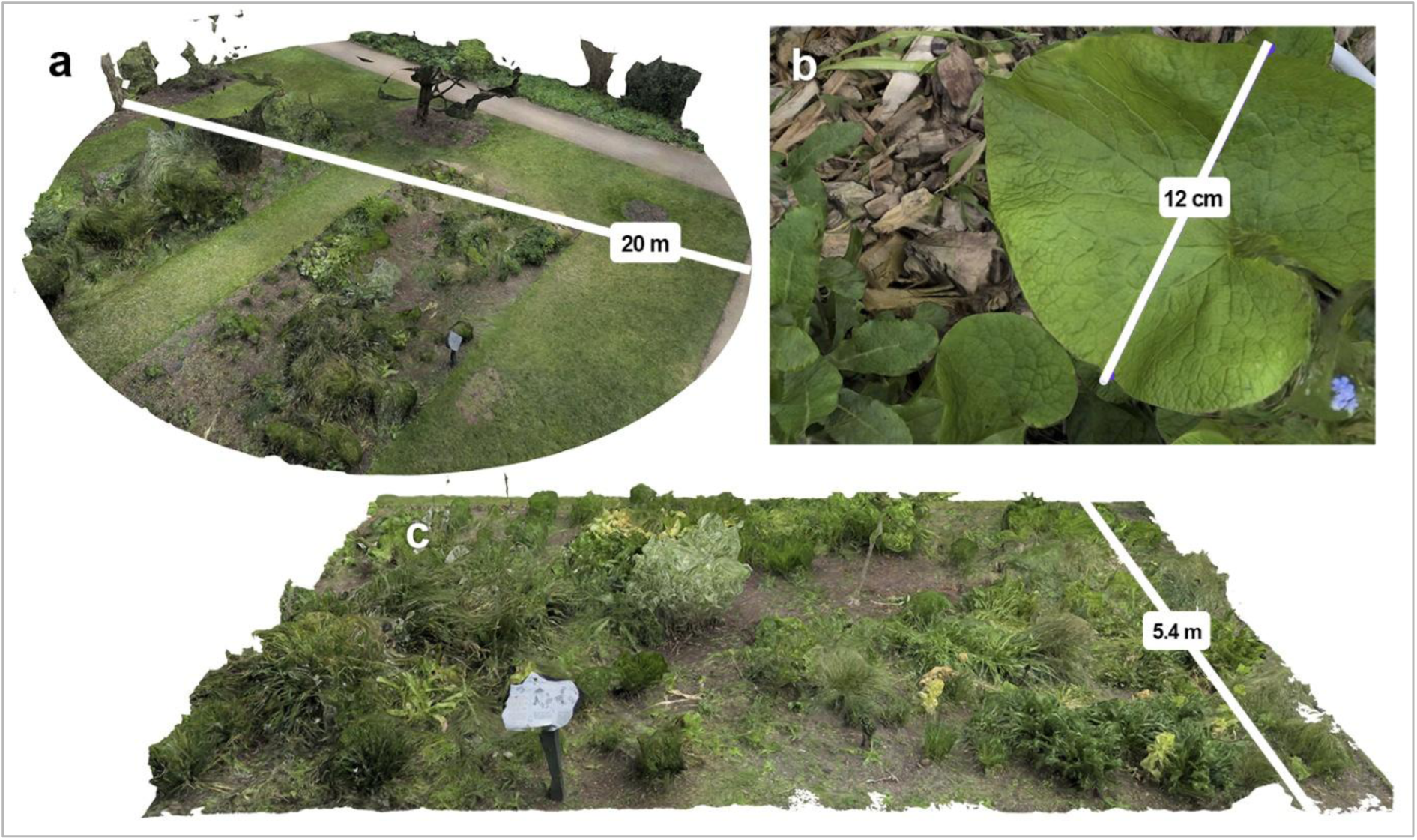
Multiple scale snapshot images from Polycam 3D mapping. **a)** Full Botanical Gardens sampling site cropped to 20 m diameter and 1.5 m height. **b)** Example of high-resolution maps achieved using RAW images **c)** 45 degree view of a single mapped flowerbed at the Botanical Gardens sampling site. Images generated by the authors via photogrammetric reconstruction using Polycam software.

To capture the micro-topography and vegetation density of the sampling sites, three-dimensional (3D) digital reconstructions were generated using photogrammetry. High-resolution RAW imagery of the plots was captured using an [Insert Camera/Phone Model, e.g., iPhone 15 Pro Max / DSLR Camera]. The image datasets were processed using the Polycam photogrammetry software platform (Polycam Inc., v2.1.19) to generate dense 3D point clouds and textured mesh models. Spatial measurements (diameters and lengths) were calibrated and verified within the software engine using known physical ground truths.

### Air sampling

Air sampling and analysis were carried out as per Kay et al. ^84^. In short, a GilAirÒ pump (Sensidyne, LP, St. Petersburg, FL, USA) collected air through 6 replicate Markes (Markes international Ltd.

Bridgend, UK) Tenax® TA tubes using a flow rate of 600ml/min at a height of 1.5 m. Sampling took place for 2 hours between the hours of 11:00 - 13:00 UST. Thermal-desorption-gas-chromatography-mass-spectrometry (TD-GC-MS) was carried out at the Jodrell laboratory, Royal Botanic Gardens, Kew using a Markes TD100-xr automated thermal desorption unit, a Tenax® TA packed cold trap (set to 4 °C) and a J&W DB-5ms GC column (30 m × 0.25 mm × 0.25 µm within an Agilent 8890 GC System (Agilent Technologies, Santa Clara, CA, USA) and an (v) Agilent 5977C GC/ MSD.

### GC-MS Library development

For identification, the spectral library previously developed by Kay et al. ^84^ was used. In short, a reduced version of NIST20 was created by retrieving all compounds present in ChEBI (release 247, December 2025) by CAS registry number.

Compounds were assigned to structural classes on the basis of molecular structure using a fixed precedence: terpene skeleton, then nitrogen-, sulphur- or halogen-containing, then aromatic ring, then oxygen-containing, then hydrocarbon. Assignments were verified independently against the ChEBI structural hierarchy (release 247, December 2025). Compounds were located in ChEBI by CAS registry number where possible and by name or synonym otherwise, as ChEBI records do not always carry a CAS number; 259 of the 262 compounds were located by one of these routes, and 207 of the 245 for which ChEBI resolves a structural class agree with the class assigned here. Remaining differences reflect the precedence ordering above, since a compound may belong to more than one structural class.

For source evidence, records were retrieved from five compound databases - NPClassifier, LOTUS, NPAtlas, KEGG and LIPID MAPS^86–90^ - of which 37 of the 262 compounds are present in at least one, reflecting their focus on natural products and metabolites. Emission source profiles were retrieved from the US EPA SPECIATE database^91^, in which 125 of the 262 compounds are listed. Natural and artificial pollution source information was retrieved from PubChem^92^, largely derived from the Hazardous Substances Data Bank; 96 compounds carry one or both, the remainder being predominantly branched alkanes and minor oxygenates not covered by these resources. Where a CAS number resolved to a stereospecific PubChem record without source data, the corresponding generic record was used. Occurrence in plant taxa was retrieved from PubChem’s consolidated taxonomy compiled from KNApSAcK, LOTUS, MetaboLights and NPASS, with organisms assigned to plants, fungi, bacteria or animals using NCBI taxonomy divisions; 226 compounds have recorded plant occurrence, including all terpenes and terpenoids.

Likely source in this setting is an interpretation for the sampling context of this study rather than a property of the compound: all sampling was conducted outdoors in vegetated greenspaces, away from indoor environments in which consumer-product emissions accumulate. Biogenic evidence was taken as a record in a curated natural-product resource - NPClassifier, LOTUS or NPAtlas - a PubChem Natural Pollution Sources entry, recorded occurrence in ten or more plant genera, or SPECIATE profiles that are predominantly vegetation-associated. Anthropogenic evidence was taken as a PubChem Artificial Pollution Sources entry, or three or more SPECIATE profiles of which at least half are in industrial, traffic, solvent or consumer-product sectors. Compounds with evidence of both kinds were annotated as mixed, except where anthropogenic profiles were strongly dominant (at least twenty profiles and at least 90 per cent of listings) and no curated natural-product record existed, in which case they were annotated as anthropogenic. Terpenes and terpenoids were mostly annotated as biogenic in this setting, with those also carrying anthropogenic evidence marked as dual-source compounds. Compounds with no evidence of either kind were annotated as undetermined.

### Analysis of VOC profiles

The reduced NIST20 library was combined with a non-isothermal Kovats retention index ^93^ for analysis in AMDIS (version 2.73) using spectral and RI matching. A minimum match factor threshold of 70 (Net), a RI window of +/- 10, and a very strong RI penalty were used, with deconvolution settings of component width of 25, an adjacent peak subtraction of two, medium shape requirements, high sensitivity, and medium resolution. Best-match compounds for each retention time and replicate tube were then chosen based on highest Net Match score. Peak areas of any compounds identified in blank control tubes were removed from the peak areas of all samples within a GC-MS run. Missing area values for compounds (i.e. those compounds found only in some technical replicate tubes) were zero-filled. The relative amount (%) of each compound - (peak area of compound / TIC of sample) × 100 - was used for all analysis. Relative amounts are therefore expressed as a percentage of the total ion signal of the sample and are directly comparable between compounds, samples and chemical classes.

### Temporal dynamics and composition of terpene and terpenoid emissions

For each site and sampling date, the summed relative abundance of all identified terpene and terpenoid was used. Additionally, compounds were classified into four functional groups - monoterpenes, monoterpenoids, sesquiterpenes, and sesquiterpenoids. Finally, to examine compound-level patterns, a heatmap was generated from mean peak areas for each compound × site × date combination. These values were min–max normalised to 0–1 using the formula (x − min) / (max − min), with a single global minimum and maximum taken across the entire matrix of compounds, sites and dates rather than compound by compound, so that colour intensity is comparable both within and between compounds. Compounds not detected in a given sample were excluded from the calculation of the global minimum and maximum and are displayed as absent rather than as zero. All analyses and visualisations were conducted in R (v.4.x) using tidyverse and ggplot2.

### Statistical modelling of site-level drivers of terpene/terpenoid abundance

To test for correlations between chemodiversity and ecological metrics, site-level terpene/terpenoid abundance was calculated by averaging peak areas across technical replicates and summing compounds to obtain total terpene/terpenoid abundance for each site × date. Values were then scaled between 0 and 1 within each sampling date to control for day-to-day variation. Response values were paired with vegetation and structural metrics derived from field surveys, including taxonomic richness (at the family, genus, and species levels), canopy cover, ground vegetation cover, tree count, and vegetation volume within multiple areas. All predictors were centred and scaled prior to analysis. To reduce multicollinearity, separate single-predictor linear mixed-effects models were fitted in R using lme4, with the form: Terpene ∼ Predictor + (1 | Location) + (1 | Date). Models were fitted by maximum likelihood, and fixed effects were assessed using Wald t-tests via lmerTest. P-values were adjusted for multiple testing using the Benjamini–Hochberg false discovery rate procedure. Model results were summarised using coefficient plots showing effect sizes ± SE and heatmaps of overall patterns.

For the species-level correlation analyses, relationships between terpene/terpenoid abundance and biodiversity metrics were assessed using ordinary linear models (lm) in R. Predictor variables were standardised where appropriate, and fitted relationships were visualised using scatterplots with fitted regression lines. Results from these models were exported for plotting and summary.

## Results

### Vegetation Structure

Vegetation structure and plant richness varied substantially among the five sampling sites (Table 1). The Botanical Gardens outdoors showed the greatest ornamental and taxonomic diversity, with five trees partially within the 20 m survey diameter and a highly managed understory dominated by ornamental and non-native herbaceous species. University Parks Dense also contained five trees, mainly horse chestnut and Austrian pine, with a ground layer of ruderal and shade-tolerant herbaceous species such as ivy, dandelion, white clover, and cow parsley. University Parks Open had no trees within the sampled diameter and supported a simpler grassland assemblage dominated by white clover, dandelion, creeping buttercup, ribwort plantain, and red sorrel. Warneford Meadow contained three trees, two oaks with one Norway spruce, and a meadow-style understory including creeping thistle, nettle, ribwort plantain, and yarrow. Wytham Woods site was adjacent to more dense woodland, yet only 5 trees were within the 20 m diameter circle from the sampling area. This site contained 2 oak, and 3 Norway spruce, and an understory characteristic of woodland edge and scrub, including bramble, hazel, honeysuckle, elder, ivy, and cleavers. Full details of plants found in quadrats are in Supplementary Table S 1.

**Table 1:** Summary of vegetation structure and biodiversity metrics across the five outdoor sampling sites at two spatial scales (20 m and 100 m diameter). Taxonomic richness (total and unique counts of families, genera, and species), vegetation structure (canopy cover, vegetation cover, and tree count), and vegetation volume estimates. “Green ground” represents the percentage cover of undeveloped ground. Vegetation volume is expressed as estimated vegetation volume (m³), with values reported for total vegetation volume, vegetation volume below 1.5 m height, and vegetation volume above 1.5 m height (tree volume).

| Metric | Survey Diameter | Oxford Botanical Garden (BO) | University Parks Dense (UD) | University Parks Open (UO) | Warneford (WA) | Wytham (WY) |
| --- | --- | --- | --- | --- | --- | --- |
| Total family in quadrats | 20 m | 20 | 13 | 6 | 10 | 18 |
| Total genus in quadrats | 20 m | 32 | 14 | 7 | 12 | 20 |
| Total species in quadrats | 20 m | 48 | 14 | 7 | 12 | 20 |
| Canopy cover (%) | 20 m | 33.52 | 91.54 | 0 | 25.21 | 31.02 |
| ‘Green’ ground (%) | 20 m | 62.68 | 94.29 | 100 | 100 | 100 |
| Tree count | 20 m | 5 | 5 | 0 | 3 | 5 |
| Vegetation volume (m<sup>3</sup>) above 1.5 m | 20 m | 551.32 | 10,252.11 | 0.00 | 968.73 | 1,609.55 |
| Vegetation volume (m<sup>3</sup>) below 1.5 m | 20 m | 81.62 | 4.75 | 1.30 | 6.77 | 47.26 |
| Vegetation volume (m<sup>3</sup>) Total | 20 m | 632.94 | 10,256.86 | 1.30 | 975.51 | 1,656.81 |
| Canopy cover (%) | 100 m | 41.16 | 31.38 | 17.25 | 6.54 | 63.5 |
| ‘Green’ ground (%) | 100 m | 46.27 | 77.91 | 89.79 | 100 | 85.58 |
| Tree Count estimate | 100 m | 45 | 42 | 12 | 12 | 80 |

Oxford Botanic Garden had the highest plant richness across all three taxonomic levels, with the greatest numbers of families, genera, and species recorded in the quadrats, whereas University Parks Open Grassy had the lowest values (Table 1). At the 20 m scale, University Parks Dense Tree area had the highest canopy cover and total vegetation volume, reflecting the presence of large, mature trees, while Oxford Botanic Garden had the greatest vegetation volume below 1.5 m, consistent with its abundant flowerbeds and lower-stature planting. As expected, University Parks Open Grassy had no trees and no canopy cover within the 20 m diameter. At the 100 m scale, canopy cover, ground vegetation cover, and estimated tree abundance continued to vary markedly among sites, with Wytham Woods showing the highest estimated tree count.

### Terpene-rich scentscapes are site-specific and temporally dynamic

Terpene and terpenoid abundance varied strongly through time and among sites (Figure 3). Across the outdoor locations, total terpene/terpenoid abundance generally increased from early spring into late spring or summer, although both the timing and magnitude of this increase differed between sites. Peak terpene/terpenoid amounts were recorded either in late spring (April 30^th^) at the Botanic Gardens, Wytham Woods, and University Parks Open, or in mid-summer (June 30^th^) at Warneford Meadow and University Parks Dense. The Botanic Garden’s site consistently showed the highest total abundance in terpenes and terpenoids followed by Wytham Woods and then Warneford Meadow. Both University Parks sites had lower overall terpene and terpenoid abundance across the sampling period. These abundance patterns were also reflected in compound counts/richness (as shown by numbers on Figure 3a).

**Figure 3:**
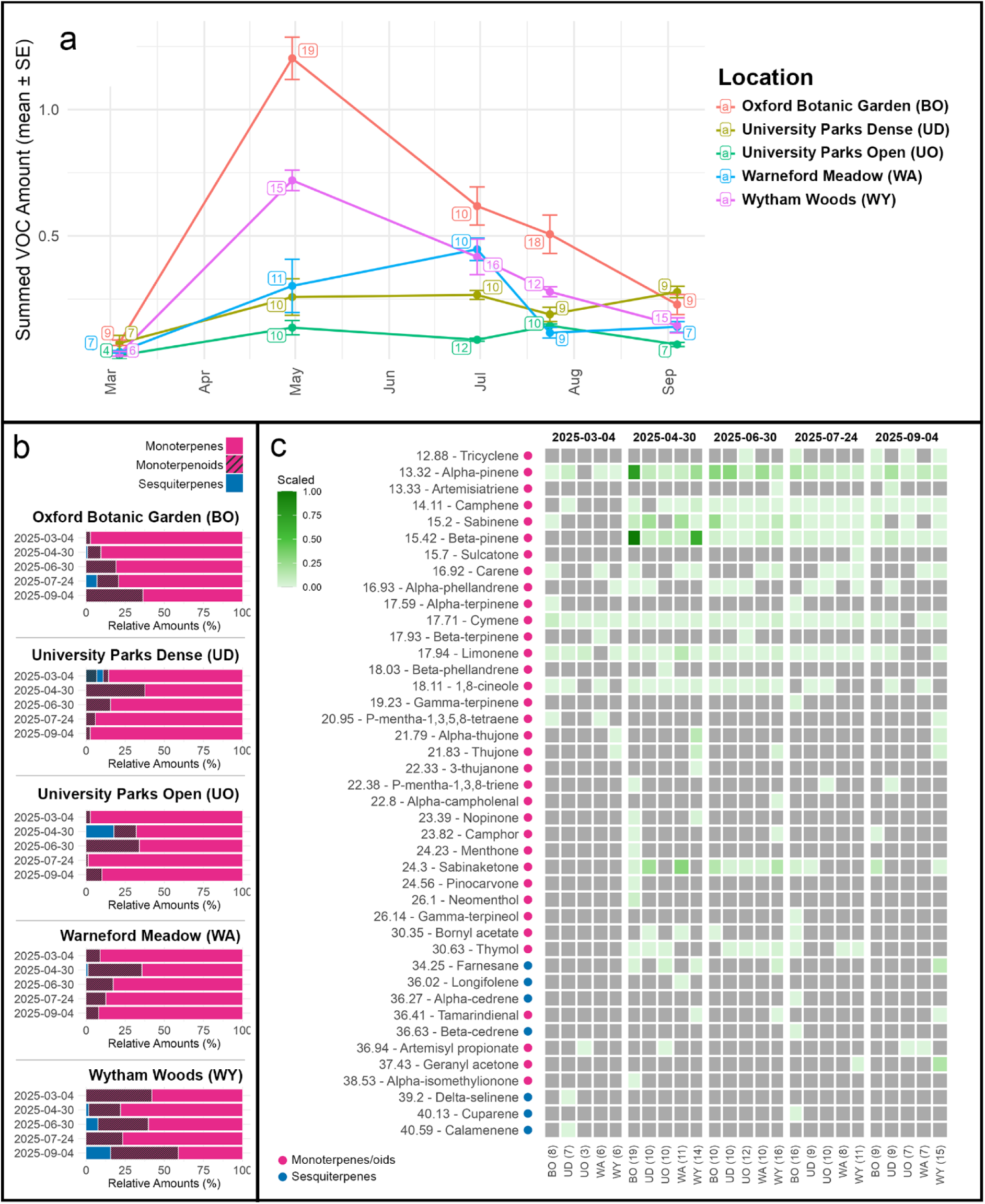
Temporal variation in terpene and terpenoid abundance and composition across five urban greenspaces. **a)** Mean (± SE) summed relative abundance of terpene- and terpenoid-class VOCs across sampling dates (March–September 2025) for Oxford Botanic Garden (BO), University Parks Dense (UD), University Parks Open (UO), Warneford Meadow (WA), and Wytham Woods (WY). Numbers adjacent to points indicate the total number of detected terpene/terpenoid compounds at each site and date. **b)** Relative contributions (%) of monoterpenes, monoterpenoids, and sesquiterpenes to the total terpene/terpenoid fraction at each site across sampling dates. **c**) Heatmap showing the scaled relative abundance of individual terpene and terpenoid compounds across sites and sampling dates. Green intensity indicates higher scaled abundance; values were min–max normalised across the entire dataset, so intensities are comparable both within and between compounds and within and between dates. Grey indicates that the compound was not detected in that sample. Compounds are grouped by class, with retention times and compound identities shown on the y-axis. Data are 4 - 6 technical replicate sampling tubes per site per timepoint.

The relative composition of the terpene/terpenoids detected was consistently dominated by monoterpenes (in all 25 out of 25 samples, Figure 3b). There was no clear seasonal or site-level pattern indicating when terpenoids became relatively more abundant. Sesquiterpenes were also only intermittently detected and showed no obvious temporal or spatial pattern. The terpene/terpenoid heatmap revealed substantial turnover in individual compounds through time and among locations (Figure 3c). Several compounds were repeatedly detected across multiple sites and dates, including α-pinene, cymene, and limonene, suggesting a shared outdoor terpene profile across the study sites. Some compounds, including camphene, sabinene, and β-pinene, became more prominent after the early spring sampling point, indicating a seasonal increase in specific monoterpenes. In contrast, 1,8-cineole (eucalyptol) appeared to diminish later in the season.

### Plant richness and the presence of mid-height vegetation are strong predictors of terpene/terpenoid presence at nose height

Plant taxonomic richness was consistently positively correlated with summed terpene/terpenoid relative abundance (Figure 4). The strongest and most consistent relationships occurring from late spring to mid-summer (April–July) when species and genus richness explained a large proportion of variation, with species richness showing the strongest positive effect (*p* < 0.01, R² = 0.795 and *p* < 0.01, R² = 0.834 respectively). By September, relationships remained positive but non-significant.

**Figure 4:**
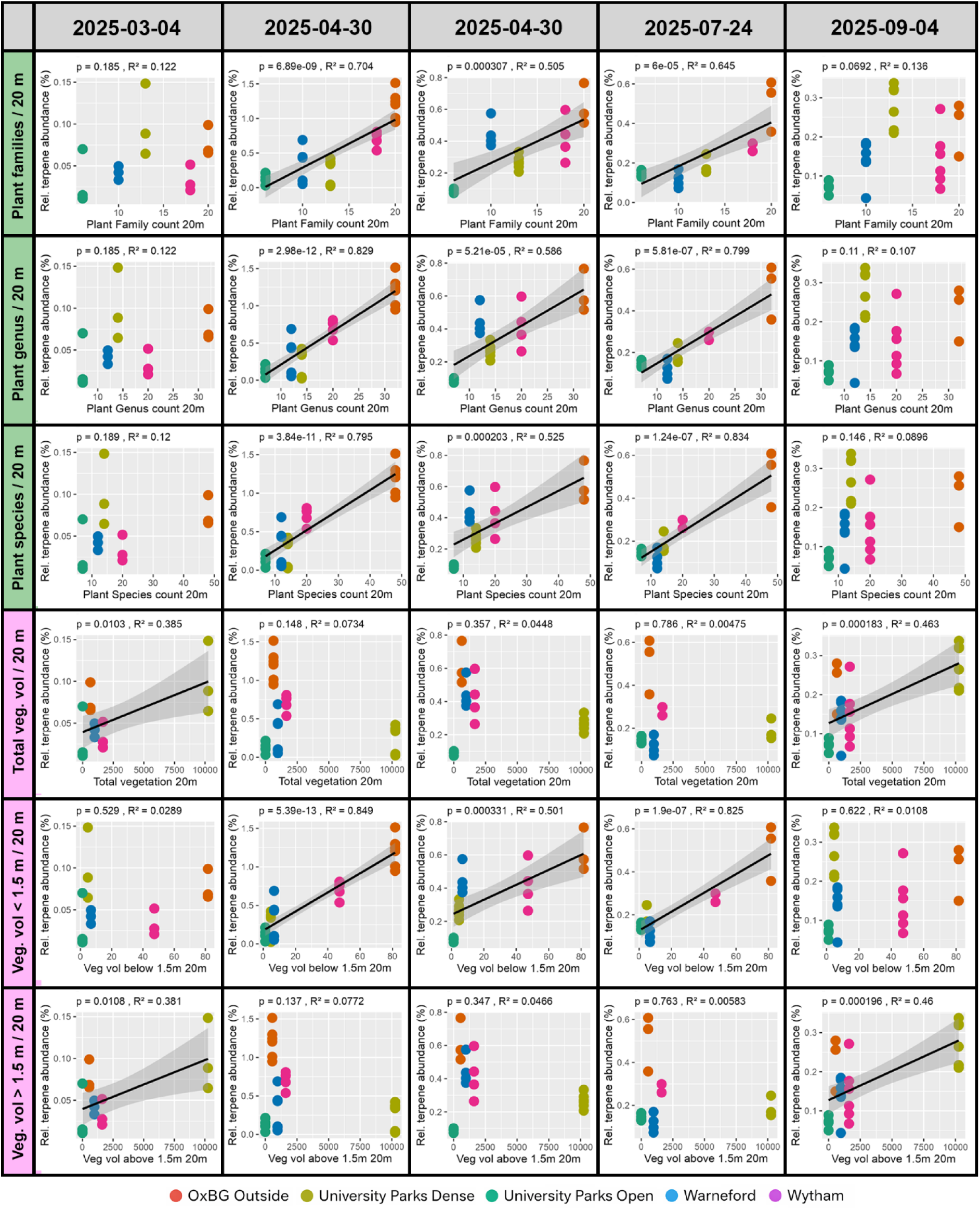
Relationships between plant taxonomic richness and vegetation volume and summed VOC relative abundance across sampling dates. Scatterplots show relationships between summed VOC relative amounts (%) and plant richness/vegetation volume within 20 m of each sampling point. Columns represent sampling dates from March to September 2025. Rows show total family richness, genus richness, and total species richness, and vegetation volume (Veg vol.) at different heights. Points represent technical replicate tubes and are coloured by location. Black lines show fitted linear regressions with grey 95 % confidence intervals and appear only when correlations are significant. Reported p-values and R² values indicate the strength of each richness–VOC relationship for each date and taxonomic level/height level. Data are 4 - 6 technical replicate sampling tubes per site per timepoint.

Across all dates, richness at finer taxonomic resolution (species and genus) was a better predictor of increased VOC abundance than family-level richness. Vegetation volume also significantly correlated with terpene/terpenoid abundance, but this depended strongly on height. Vegetation volume within the 0 – 1.5 m layer showed the strongest and most consistent positive associations with significant relationships (*p* < 0.001) on April 30^th^, June 30^th^, and July 24^th^ (R² = 0.849, 0.501, and 0.825 respectively). In contrast, total vegetation volume across the full 20 m plot was a weaker and less consistent predictor, with significance only in March (*p* = 0.01, R² = 0.392) and September (*p* < 0.001, R² = 0.453). Overall, summed VOC abundance was most strongly associated with local plant diversity and vegetation volume within the lower-to-mid vegetation layer.

### Tree abundance, canopy cover, and the presence of anthropogenic structures had fewer positive correlations with terpene/terpenoid abundance

Overall, terpene/terpenoid abundance showed a similar number of positive correlations with tree abundance, canopy cover, and the presence of anthropogenic structures within a 100 m diameter and a 20 m diameter (Figure 5). For example, local canopy cover within 20 m was significant in March and September only, with moderate explanatory power in March (*p* < 0.01, R² = 0.491) and September (*p* < 0.01, R² = 0.578), whereas canopy cover at 100 m showed significant positive relationships in April (*p* < 0.01, R² = 0.328), and July (*p* < 0.01, R² = 0.398). Tree counts showed consistent positive associations within 20 m diameter, although the R² values were consistently weak. In contrast, tree count within 100 m diameter significantly correlated with terpene/terpenoid abundance only in April (*p* = 0.003, R² = 0.27) and July (*p* = 0.016, R² = 0.312). Green ground cover – the percentage of ground absent of anthropogenic development such as paths and buildings – showed strong and consistent negative relationships with VOC abundance at both 20 m and 100 m scales.

**Figure 5:**
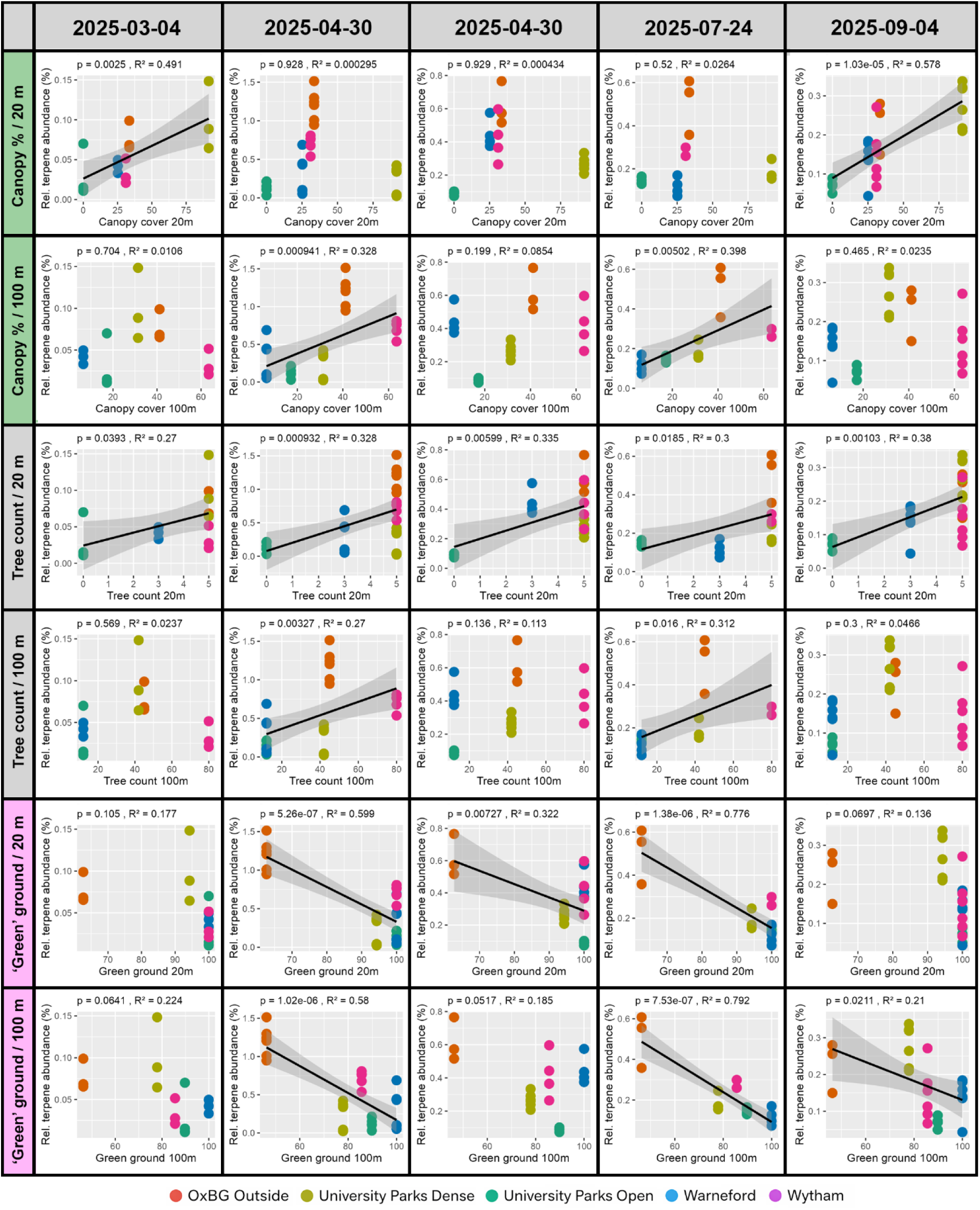
Scatterplots show relationships between summed VOC relative amounts (%) and vegetation structure variables measured around each sampling point. Columns represent sampling dates from March to September 2025. Rows show canopy cover (%), tree count, and ground “green” cover (%) within 20 m and 100 m. Points represent technical replicate tubes and are coloured by location. Black lines show fitted linear regressions with grey 95 % confidence intervals and appear only when correlations are significant. Reported p-values and R² values indicate the strength of each vegetation structure–VOC relationship for each date and spatial scale. Data are 4 - 6 technical replicate sampling tubes per site per timepoint.

### Plant richness and vegetation height are the strongest predictors of plant based volatile abundance, but weak predictors of anthropogenic volatile abundance

To assess whether the significant positive relationships between vegetation characteristics and terpene/terpenoid abundance were specific to terpenes or reflected broader patterns across both plant-derived and anthropogenic volatiles, we extended the analysis to other major VOC chemical classes, including aliphatics, aromatics, oxygenated aliphatics, and other heteroatom-containing compounds. For each VOC class, p-values from the single-predictor models were adjusted using the Benjamini-Hochberg FDR procedure, and adjusted p-values were used to denote significance. Across chemical classes, the strength and direction of relationships varied substantially (Figure 6). Terpenes, as shown above, showed the strongest and most consistent positive associations with vegetation characteristics, particularly with plant genus richness (*p* < 0.001) and mid-height vegetation volume (*p* = 0.04). Oxygenated aliphatics, which include a range of plant-associated and secondary biogenic compounds, exhibited a similar pattern, with positive associations with plant family and genus richness (*p* = 0.01, 0.03 respectively) and vegetation volume below 1.5 m (*p* = 0.02). Only plant genus became non-significant after multiple-testing correction (*p* = 0.09). In contrast, aliphatics (*p* > 0.45 in all tests) and aromatics (*p* > 0.92 in all tests) showed no significant correlations to terpene/terpenoid abundance. The “other” compound class (N-, S-, and halogen-containing compounds) also showed limited and inconsistent associations (*p* > 0.36 in all tests).

**Figure 6:**
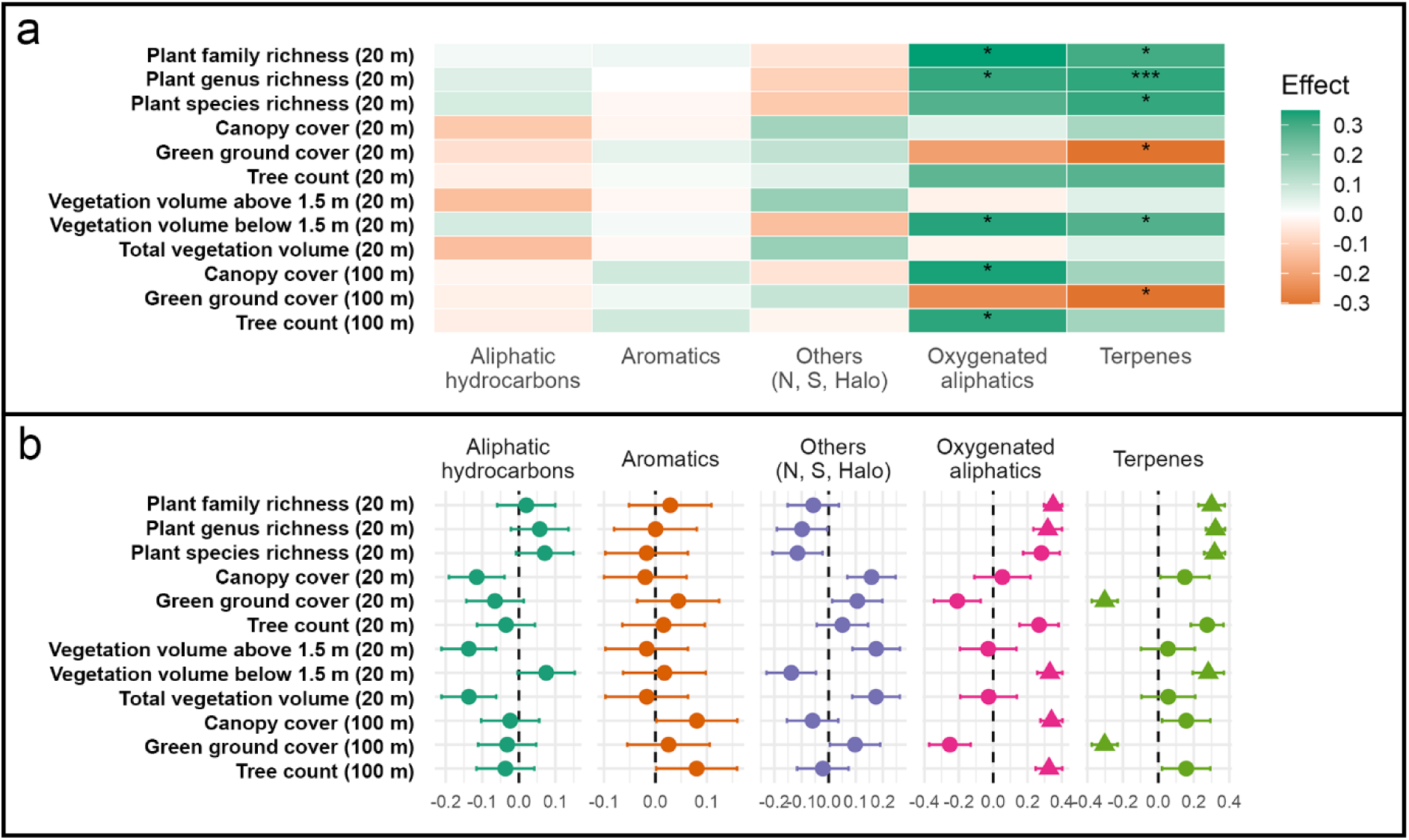
Relationship between vegetation structure, biodiversity, and VOC chemical classes. **a**) Heatmap showing standardised effect sizes from single-predictor linear mixed-effects models relating vegetation metrics to summed VOC abundance for each chemical class. VOCs were grouped into five classes: aliphatic hydrocarbons, aromatics, other compounds (N-, S-, and halogen-containing), oxygenated aliphatics, and terpenes (including terpenoids). Colours indicate effect size, with green representing positive relationships and orange representing negative relationships. Stars denote FDR-adjusted significance (* p < 0.05, ** p < 0.01, *** p < 0.001). Models included random intercepts for location and sampling date. **b**) Forest plots showing standardized effect sizes from the single-predictor linear mixed-effects models. The dashed vertical line marks zero effect, and marker shape denotes significance category (triangles show significant results). Data are 4 - 6 technical replicate sampling tubes per site per timepoint. Counts per class are terpenes 22, terpenoids 20, oxygenated aliphatics 64, aromatics 72, aliphatic hydrocarbons 57, others 27.

## Discussion

Despite increasing interest in plant-derived volatile organic compounds (VOCs) as contributors to human wellbeing, it remains unclear which features of greenspaces most affect the distribution and availability of health-promoting compounds ^46^. This study aimed to address this question by assessing biogenic VOC presence and relative abundance across five contrasting greenspace types in Oxford, UK, and correlating these patterns to multiple biodiversity and vegetation structure metrics.

Plants are known to be major sources of biogenic volatile organic compounds (bVOCs), particularly terpenes/terpenoid ^47,48^, which are constitutively produced and tend to increase with vegetation volume due to greater foliar surface area ^49,50^. Biotic stresses such as herbivory ^51^, together with abiotic stresses such as drought, heat and high UV ^52–54^, can further influence terpene/terpenoid biosynthesis and emission ^55^. Based on this, one might expect greater terpene abundance in sites with higher total vegetation volume. However, this pattern was not observed in this work. Within the sites surveyed, University Parks Dense contains the most trees, the most canopy cover, and the largest total vegetation volume, yet it was consistently low in terpene/terpenoid abundance and diversity compared to sites such as Wytham and the Botanical Gardens.

This result could be explained using two physical characteristics of tree-based landscapes. Firstly, air moving through taller vegetation, such as urban tree canopies, has been shown to pass through more rapidly that lower vegetation due to wind-based vertical transport processes within the canopy ^45,56^. Additionally, previous work in subtropical forest systems has shown that terpene/terpenoid emission rates in understory vegetation is diminished due to the reduction in light availability and thus decreased photosynthetic activity, heat stress and drought stress ^53,57,58^. To assess this further, vegetation volume was also assessed at a height below 1.5 m to correlate with human nose height. Our results showed a significant correlation between terpene/terpenoid abundance and vegetation at this scale – a relationship led by the high level of shrubs and flowerbeds at the Botanical Gardens and dense woodland border vegetation at Wytham Woods.

The idea of mid-level vegetation has been explored by Lygum et al.^23^ who suggested tailoring plant locations and heights depending on plant species-specific factors such as fragrant leaves at foot height amongst stepping stones - this paper goes further by suggesting larger scale urban design features. Choosing which specific plants to include in urban greenspaces was not explored here, however, the tailoring of greenspace plants should be an important part of future work, for example, (i) which plants have desired levels of terpene/terpenoid emissions, high floral abundance, and elongated flowering periods (ii) which plants are known to produce inflammatory or allergic reactions; (ii) which plants are best suited to local, individual, cultural, and demographic differences^59–61^, and importantly (iv) which plants best provide the physiological or psychological effects desired in the space being used ^44^.

A 2025 paper ^18^ using photographs of urban flower meadows showed that positive emotions were evoked in participants of all ages, genders and demographic groups. The addition of the range of scents that would be produced by such a species-rich herbaceous community would likely compound these results. A rise in low-story vegetation planting such as meadows in urban areas may therefore both save costs on greenspace management, and more importantly, provide ecosystems for wildlife such as insects. Urban planting strategies such as these would therefore align well with the WHO One Health framework, which recognises that the health of humans, domestic and wild animals, plants, and the wider environment are closely linked and interdependent.

Increasing reliance on lower story vegetation would bring with it an increase in plant diversity. More diverse plant communities contain a greater number of terpene-producing taxa, each likely to contribute distinct compounds to the local volatile profile. From this, it could be hypothesised that higher plant diversity would bring with it higher levels of terpenes/terpenoids in the ambient air.

Indeed, Medina-van Berkum et al. ^35^ demonstrated that in experimental grassland systems that both the abundance and diversity of emitted terpenes/terpenoids increase with plant species richness.

Similarly, Carlson et al. ^62^ found that VOC β-diversity increased significantly in heterogeneous forests compared to more homogeneous systems. Our results support the findings of these studies by showing that terpene/terpenoid profiles are significantly positively correlated with taxonomic richness across family, genus, and species levels.

The effects of biodiversity on VOC emissions are context-dependent and will vary with many characteristics of an urban greenspace. As such, the positive relationship between plant diversity and VOC emissions shown here has not been shown universally. Some studies report reduced emissions in more diverse systems, potentially due to competitive interactions, resource limitation, or altered plant physiology ^57^. As already mentioned, terpenes play key roles in plant defence and signalling - their production is often induced (or modified) in response to biotic stressors such as herbivory and pathogen attack. More diverse plant communities have shown decreased herbivory and pathogen damage ^63,64^ and as such the contribution of adding additional terpene-producing plant taxa at high richness may be offset by reduced induction. There may therefore be a level of taxonomic richness beyond which terpene/terpenoid production ceases to rise, although our design cannot test this.

Extending our analysis across other VOC classes to include anthropogenic compounds revealed that the observed relationships between VOC presence and biodiversity metrics were not universal.

Strong correlations were identified for terpenes/terpenoids and oxygenated aliphatics, whereas relationships for other classes (aromatics, aliphatics, and others) were weak or absent. These differences correspond with the likely origins of the compounds concerned. Source annotations compiled independently of the air measurements, from curated natural-product resources, PubChem source records and US EPA SPECIATE emission profiles (Supplementary Tables S2 and S3), assigned every terpene and terpenoid with biogenic evidence, and none with anthropogenic evidence alone. The same ordering was apparent across the other classes: 31 % of oxygenated aliphatics were assigned biogenic evidence with no anthropogenic evidence, compared with 11 % of aromatics, 10 % of heteroatom-containing compounds and 6 % of aliphatic hydrocarbons, among which anthropogenic evidence predominated. Several of the terpenes detected here do have well-documented anthropogenic sources: limonene is reported predominantly from consumer-product emission profiles, and cymene and farnesane most frequently from vehicle-related profiles.

Fragranced consumer products are themselves dominated by monoterpenes rather than by other chemical classes, so the overlap between our detections and such products is expected and cannot by itself distinguish the two sources. We therefore compared our detections against a published inventory of the volatile compounds emitted by 134 common consumer products^65^. Of the 338 compounds reported here, only 37 consumer listed compounds were detected in our samples. These were almost exclusively monoterpenes that are also major constituents of plant emissions, including limonene, eucalyptol, gamma-terpinene, alpha- and beta-pinene, camphene and sabinene.

Compounds characteristic of fragranced products but not of vegetation were absent despite being represented in our spectral library: benzyl acetate (reported from 22 of the 134 products), 4-tert- butylcyclohexyl acetate (25), hexyl acetate (15), ethyl 2-methylbutyrate (12), benzyl alcohol (19) and dihydromyrcenol (27) were not detected in any of the 111 samples. Three terpenes that are common in consumer products were also absent, namely beta-trans-ocimene (49), linalool (45) and beta-myrcene (42), indicating that the terpene profile recorded here is not that of a consumer-product source. Our results therefore support the interpretation that the patterns observed in this study are driven by biological processes rather than simply reflecting overall increases in volatile concentrations in more vegetated environments due to physical factors such as microclimate.

Tree-dominated greenspaces also play an important role in propagating negative atmospheric chemistry which can lead to negative health outcomes such as respiratory and cardiovascular disease ^66^. This occurs when biogenic volatile organic compounds, particularly terpenes emitted by trees, react with other pollutants present in the urban atmosphere. In cities, these reactions are often intensified by traffic-related emissions which provide the nitrogen oxides and other reactive compounds needed to drive secondary pollutant formation. For example, tree-emitted terpenes are oxidised in the atmosphere by OH radicals, ozone, and nitrate radicals. Those oxidation products (secondary organic aerosols) are often less volatile than the original parts, so they can either condense onto existing particles or form new particles. That increases fine particulate matter, including PM2.5. Terpenes also participate in photochemical ozone formation, especially where traffic emissions supply abundant NO_x_. Estimates suggest that approximately 13 % of summertime ozone may be attributable to urban vegetation emissions ^67^.

To compound this, tree canopies can obstruct the dispersal of VOCs. While this would be of benefit for walking in traffic-free natural environments, trees next to roads in urban areas can obstruct high levels of traffic-related pollutants. Reducing their dispersion, therefore, is not desired ^68–70^. Modelling studies suggest that this aerodynamic effect can increase NO₂ concentrations by up to ∼37 % locally^71^. Dense hedges, by contrast, can act as physical barriers, reducing the transport of traffic-related pollutants to pedestrians and lowering near-road exposure by up to 15–60 % depending on their design and placement ^72–74^. Grasses and low-story flowering plants, however, have minimal impact on airflow-based pollutant trapping, while still offering modest benefits in terms of cooling and pollutant deposition ^72,75^.

This paper is not arguing for tree removal or a reduction in the planting of trees in urban greenspaces - trees have many well-established environmental benefits such as shading, the reduction of the urban heat island effect, and their contributions to stormwater management ^76–79^. Trees also remove airborne pollutants through deposition onto leaf surfaces ^71^. They are additionally highly valued by the public for their aesthetic and cultural benefits ^80^. It may be prudent, however, to prioritise trees which have been found to produce lower terpene emissions alongside low-story vegetation ^40,81^.

This study focused on the spring and summer seasons when outdoor recreation is at its highest. Consistent biodiversity metrics across all time points were used. While this could be considered a limitation as greenspaces change over time, in practice sites tend to remain relatively distinct in their overall ecological composition. This approach therefore allows us to provide evidence that even across timescales, areas of greater plant diversity (particularly where vegetation volume occurs at or below nose height) are associated with increased concentrations of plant-derived compounds available for human inhalation.

It must also be noted that this study was limited by the small number of independent sampling sites, which restricted the use of a full multivariable model. Although repeated sampling improved measurement resolution, the site-level habitat variables were still only available across five greenspaces, so the results should be interpreted as exploratory. Future studies should increase the number of sites sampled to allow stronger modelling of how multiple vegetation and habitat metrics jointly explain terpene and terpenoid variation. Additionally, volatile profiles over both longer and shorter timescales would strengthen our conclusions. Microenvironmental variables were not measured at the sampling points. Temperature, light availability and humidity influence terpene biosynthesis and emission, and wind speed and direction influence dilution and transport, so the between-date variation we report is likely to reflect these factors alongside seasonal changes in the vegetation itself. All five sites were, however, sampled simultaneously on each date and therefore under common prevailing conditions, so differences between sites are less likely to be attributable to them.

The attribution of individual compounds to biogenic or anthropogenic sources is a major difficulty for this and comparable studies. As discussed, many of the compounds detected here have both vegetation/biogenic and anthropogenic origins: limonene, α-pinene and cymene are emitted by plants but are also major constituents of cleaning products, solvents, fragrances, and personal care products. As such greenspace users represent a plausible source. We did not attempt a tracer-based source apportionment, and the origin annotations in Supplementary Table S2 should therefore be read as indicative rather than definitive. Nonetheless, three features of our data argue against a substantial anthropogenic contribution to the relationships reported here. First, the associations were class-specific: terpenes/terpenoids and oxygenated aliphatics correlated strongly with plant richness and lower-storey vegetation volume, whereas aromatics, aliphatic hydrocarbons and heteroatom-containing compounds showed no significant relationships in any test. Because fragranced consumer products emit aromatic esters and aldehydes alongside terpenes ^65,82^, a confound scaling with human presence would not be expected to respect structural class boundaries in this way. Second, the one compound in our dataset attributable specifically to fragranced products, alpha-isomethylionone, was identified in only five of 111 samples and exclusively at the Botanic Garden, at a mean relative abundance approximately 40-fold lower than that of α-pinene.

Third, monoterpenes dominated the terpene/terpenoid fraction in all 25 site × date samples, a composition characteristic of vegetation emission rather than consumer product use. Establishing source attribution with confidence will always be difficult in untargeted environmental GCMS studies such as this, however, future work could include complementary approaches such as including targeted fragrance tracers.

In conclusion, our study supports the hypothesis that enhancing human exposure to beneficial plant-derived volatiles in urban greenspaces may be better served by prioritising structurally diverse vegetation within the lower-to-mid height range rather than focusing on tree cover or total vegetation volume. Positive correlations between terpene/terpenoid abundance and plant diversity appear despite the higher percentage of anthropogenic development within the Botanical gardens sampling site. Green ground cover showed strong and consistent negative relationships with terpene/terpenoid abundance at both 20 m and 100 m scales - a result that strengthens our conclusion that more diverse greenspaces, even when in a truly urban environment, will enhance exposure to plant-derived volatiles.

It must be noted that plant-derived volatiles are not uniformly positive/benign. Terpenes can act as respiratory irritants at elevated concentrations, and the hydroperoxides formed when limonene and linalool autoxidise on exposure to air are established contact allergens ^83^. Sensitisation to these compounds has been characterised principally through skin contact with fragranced products rather than through ambient exposure outdoors, but it reinforces the case made above for selecting species deliberately rather than maximising volatile emission indiscriminately.

## Data availability

All data supporting the findings of this study are openly available from Zenodo at https://doi.org/10.5281/zenodo.22672170. The deposit contains the complete project directory as used in the analysis: the raw AMDIS deconvolution reports, the sample metadata, and the processed compound tables produced by the pipeline. The full list of compounds detected, with their classification and source annotation, is also given in Supplementary Table S2.

## Code availability

All custom code used to process the chromatographic data and to generate the figures and statistical results reported here is openly available from Zenodo at https://doi.org/10.5281/zenodo.22672170. The deposit comprises the processing pipeline, which converts the raw AMDIS deconvolution reports into the analysis-ready compound tables, and five R scripts reproducing Figures 3–6, including the linear mixed-effects models of emission drivers. The archive preserves the original directory structure, so the scripts run as deposited without modification of file paths; each script documents its inputs, outputs and method. Analyses were performed in R.

## Acknowledgements

We would firstly like to thank the University of Oxford Botanic Garden and Wytham Woods for permission to sample on their sites. We would also like to thank Geoffrey Kite of the Jodrell Laboratory (Royal Botanic Gardens, Kew), all members of the Long-Term Ecology Lab (LTEL, University of Oxford), and all researchers at the Leverhulme Centre for Nature Recovery (LCNR, University of Oxford) for their assistance and input.

## Funding

This work was funded by the Leverhulme Centre for Nature Recovery, made possible thanks to the generous support of the Leverhulme Trust - Grant number RC-2021-76.

## Author Contributions

WK developed methodologies, collected and processed the data, wrote the main manuscript text, and prepared all figures.

KW conceptualised the research, acquired funding, and edited the main manuscript text. KS and MT helped collect the field data.

All authors reviewed the manuscript.

## Competing Interests

The authors declare no competing financial or non-financial interests.

## Notes

### Competing Interest Statement

The authors have declared no competing interest.

