## Supplementary Information for "The scents of summer: Plant-diversity and vegetation-height drive health-relevant scent exposure in urban greenspaces"

### npj Clean Air

1. William T. Kay [a,b,*]
2. Kieran E. Storer [a]
3. Molly Tucker [a]
4. Katherine Willis [a]

[a] Department of Biology, University of Oxford, South Parks Road, Oxford OX1 3RB, UK

[b] Leverhulme Centre for Nature Recovery (LCNR), University of Oxford, Oxford, UK

### Supplementary Table S1

**Vegetation surveys carried out at five sampling sites using quadrats. Botanical Gardens Outdoors (BO), University Parks Dense (UD), University Parks Open (UO), Warneford Meadow (WA), Wytham Woods (WY).**

| | |
|---|---|
| BO | *Cordyline australis* |
| BO | *Galanthus galatea* |
| BO | *Hamamelis 'Orange Peel'* |
| BO | *Junpiperus chinensis* |
| BO | *Laurus azorica* |
| BO | *Liriodendron chinense* |
| BO | *Magnolia doltsopa* |
| BO | *Juniperus chinensis* |
| BO | *Allium flavum* |
| BO | *Allium karataviense* |
| BO | *Allium oreophilum* |
| BO | *Allium rosenbachianum* |
| BO | *Allium siculum* |
| BO | *Allium sphaerocephalum* |
| BO | *Aristolochia tomentosa* |
| BO | *Aster divaricatus* |
| BO | *Bletilla striata 'Albo-striata'* |
| BO | *Camassia cusickii* |
| BO | *Chamaerops humilis* |
| BO | *Cortaderia selloana* |
| BO | *Crocus sieberi 'Firefly'* |
| BO | *Crocus speciosus* |
| BO | *Fargesia scabrida* |
| BO | *Galium odoratum* |
| BO | *Geranium sylvaticum 'Mayflower'* |
| BO | *Helleborus orientalis 'Ballard''* |
| BO | *Iris danfordiae* |
| BO | *Kniphofia citrina* |
| BO | *Lilium martagon 'Album'* |
| BO | *Lilium regale* |
| BO | *Lilium regale* |
| BO | *Narcissus Ã - incomparabilis* |
| BO | *Narcissus lobularis* |
| BO | *Narcissus poeticus var. recurvus* |
| BO | *Nassella trichotoma* |
| BO | *Sauromatum venustum* |
| BO | *Scilla bifolia* |
| BO | *Scilla mischtschenkoana* |
| BO | *Scilla peruviana* |
| BO | *Sternbergia lutea* |
| BO | *Tulipa kolpakowskiana* |
| BO | *Tulipa turkestanica* |
| BO | *Yucca gloriosa 'Variegata'* |
| BO | *Zantedeschia albomaculata* |
| UD | *Taraxacum officinale* |
| UD | *Hedera helix* |
| UD | *Petroselinum crispum* |
| UD | *Trillium chloropetalum* |
| UD | *Iris foetidissima* |
| UD | *Trifolium repens* |

| | |
|---|---|
| UD | *Bellis perennis* |
| UD | *Geum urbanum* |
| UD | *Ranunculus repens* |
| UD | *Primula hendersonii* |
| UD | *Glechoma hederacea* |
| UD | *Viburnum rhytidophyllum* |
| UD | *Aesculus hippocastanum* |
| UD | *Pinus sylvestris* |
| UO | *Trifolium repens* |
| UO | *Taraxacum officinale* |
| UO | *Ranunculus repens* |
| UO | *Rumex acetosa* |
| UO | *Bellis perennis* |
| UO | *Stellaria media* |
| UO | *Plantago lanceolata* |
| WA | *Cirsium arvense* |
| WA | *Rumex acetosa* |
| WA | *Urtica dioica* |
| WA | *Plantago lanceolata* |
| WA | *Taraxacum officinale* |
| WA | *Heracleum phondylium* |
| WA | *Cerastium fontanum* |
| WA | *Trifloum repens* |
| WA | *Ranunculus repens* |
| WA | *Jacobaea vulgaris* |
| WA | *Quercus robur* |
| WA | *Quercus robur* |
| WA | *Picea abies* |
| WY | *Rubus fruticosus* |
| WY | *Corylus avellana* |
| WY | *Lonicera periclymenum* |
| WY | *Sambucus nigra* |
| WY | *Hedera helix* |
| WY | *Urtica dioica* |
| WY | *Glechoma hederacea* |
| WY | *Nepeta cataria* |
| WY | *Trifolium repens* |
| WY | *Taraxacum officinale* |
| WY | *Ranunculus repens* |
| WY | *Plantago major* |
| WY | *Rumex acetosa* |
| WY | *Heracleum sphondylium* |
| WY | *Petroselinum crispum* |
| WY | *Frangula alnus* |
| WY | *Polygonatum multiflorum* |
| WY | *Rosa canina* |
| WY | *Quercus robur* |
| WY | *Picea abies* |

### Supplementary Table S2

Chemical classification and source evidence for all 262 compounds detected across the five outdoor sampling sites. Columns give the structural class assigned in this study, with the class derived independently from ChEBI in parentheses where the two differ; the compound databases holding a record for each compound; evidence from US EPA SPECIATE emission source profiles; the PubChem record from which source information was retrieved and whether it contains natural or artificial pollution source information; our interpretation of the likely source in this sampling context; and recorded occurrence in plant taxa with example genera.

| CAS | Name | Chemical class (ChEBI-derived class) | Found in | SPECIATE evidence | PubChem CID used | Natural source (PubChem) | Artificial source (PubChem) | Likely source in this setting | Plant occ. | Plant genera (examples) |
|---|---|---|---|---|---|---|---|---|---|---|
| 110-93-0 | Sulcatone | Terpenoids (Oxygenated aliphatics) | NPClassifier, NPAtlas, LIPID MAPS | Not listed in SPECIATE | 9862 | Yes | No | Biogenic | Yes | *Durio; Lablab; Helianthus; Vaccinium; Opuntia; Ocimum; Artocarpus; Capsicum (+72 more)* |
| 16982-00-6 | Cuparene | Terpenes (Terpenoids) | NPClassifier, LIPID MAPS | Not listed in SPECIATE | 86895 | No | No | Biogenic | Yes | *Frullania; Isodon; Linaria; Capsicum; Portulaca; Solanum; Morella; Phaseolus (+78 more)* |
| 28624-28-4 | Delta-selinene | Terpenes | NPClassifier, LIPID MAPS | Not listed in SPECIATE | 12308846 | No | No | Biogenic | Yes | *Vaccinium; Averrhoa; Phaseolus; Prunus* |
| 475-20-7 | Longifolene | Terpenes | NPClassifier, KEGG, LIPID MAPS | Not listed in SPECIATE | 1796220 | No | No | Biogenic | Yes | *Artemisia; Gastrodia; Oryza; Capsella; Sicyos; Glycyrrhiza; Tripterygium; Dalbergia (+54 more)* |
| 99-83-2 | Alpha-phellandrene | Terpenes | NPClassifier, KEGG, LIPID MAPS | Not listed in SPECIATE | 7460 | Yes | No | Biogenic | Yes | *Prunus; Asparagus; Barbilophozia; Pimenta; Castanea; Mammea;* |

| | | | | | | | | | | |
|---|---|---|---|---|---|---|---|---|---|---|
| | | | | | | | | | | *Dioscorea; Brassica (+72 more)* |
| 99-86-5 | Alpha-terpinene | Terpenes | NPClassifier, KEGG, LIPID MAPS | Not listed in SPECIATE | 7462 | No | No | Biogenic | Yes | *Lablab; Helianthus; Vaccinium; Opuntia; Morus; Rubus; Sambucus; Vigna (+77 more)* |
| 498-15-7 | Carene | Terpenes | NPClassifier, KEGG, LIPID MAPS | Not listed in SPECIATE | 443156 | No | No | Biogenic | Yes | *Empetrum; Fragaria; Glycine; Mentha; Origanum; Rubus; Triticum; Vaccinium (+91 more)* |
| 99-87-6 | Cymene | Terpenes | NPClassifier, KEGG, LIPID MAPS | 664 profiles (8 biogenic); top sectors: Mobile (428); Mobile; Onroad (104); Mobile; Onroad; Light Duty (92) | 7463 | Yes | Yes | Biogenic (dual-source compound) | Yes | *Toona; Wisteria; Fumaria; Lablab; Byrsonima; Aglaia; Asparagus; Juglans (+76 more)* |
| 99-85-4 | Gamma-terpinene | Terpenes | NPClassifier, KEGG, LIPID MAPS | 11 profiles (7 biogenic); top sectors: Biomass Burning; Prescribed Fire (4); Biomass Burning; Wildfire (4); Pulp And Paper (3) | 7461 | No | No | Biogenic | Yes | *Piper; Cerbera; Eucalyptus; Juglans; Mentha; Rubus; Sambucus; Vaccinium (+76 more)* |
| 5989-27-5 | Limonene | Terpenes | NPClassifier, KEGG, LIPID MAPS | 137 profiles (7 biogenic); top sectors: Consumer Products (93); Degreasing (7); Volatile Chemical Products (6) | 440917 | Yes | Yes | Biogenic (dual-source compound) | Yes | *Pinellia; Punica; Virola; Bertholletia; Citrus; Lotus; Brasenia; Solanum (+75 more)* |
| 508-32-7 | Tricyclene | Terpenes | NPClassifier, LIPID MAPS | Not listed in SPECIATE | 79035 | No | No | Biogenic | Yes | *Capsicum; Artemisia; Cocos; Attalea; Eruca; Musa; Carum; Cotoneaster (+57 more)* |

| | | | | | | | | | | |
|---|---|---|---|---|---|---|---|---|---|---|
| 471-15-8 | 3-thujanone | Terpenoids | NPClassifier, NPAtlas, LIPID MAPS | Not listed in SPECIATE | 91456 | No | No | Biogenic | Yes | *Curcuma; Diospyros; Liatris; Arabidopsis; Pseudopodospermum; Funtumia; Hansenia; Cirsium (+43 more)* |
| 546-80-5 | Alpha-thujone | Terpenoids | NPClassifier, KEGG, LIPID MAPS | Not listed in SPECIATE | 261491 | No | No | Biogenic | Yes | *Laurus; Atriplex; Carthamus; Averrhoa; Allium; Castanea; Brassica; Nicotiana (+59 more)* |
| 5655-61-8 | Bornyl acetate | Terpenoids | NPClassifier, KEGG, LIPID MAPS | 1 profiles; top sectors: Composting (1) | 93009 | No | No | Biogenic | Yes | *Paraleonurus; Strychnos; Pteris; Capsicum; Linum; Ocimum; Distylium; Calea (+81 more)* |
| 76-22-2 | Camphor | Terpenoids | NPClassifier, LOTUS, KEGG, LIPID MAPS | 44 profiles (2 biogenic); top sectors: Biomass Burning; Wildfire (18); Consumer Products (17); Pulp And Paper (3) | 2537 | Yes | Yes | Biogenic (dual-source compound) | Yes | *Vaccinium; Ribes; Theobroma; Amelanchier; Capsicum; Solanum; Portulaca; Lupinus (+77 more)* |
| 1196-31-2 | Menthone | Terpenoids | NPClassifier, KEGG, LIPID MAPS | Not listed in SPECIATE | 26447 | Yes | Yes | Biogenic (dual-source compound) | Yes | *Gymnema; Capsicum; Codonopsis; Petasites; Solanum; Veratrum; Phlebodium; Gentiana (+74 more)* |
| 2216-52-6 | Neomenthol | Terpenoids | NPClassifier, KEGG, LIPID MAPS | Not listed in SPECIATE | 1254 | Yes | Yes | Biogenic (dual-source compound) | Yes | *Tephroseris; Brickelliastrum; Ricinus; Brassica; Luffa; Vigna; Vaccinium; Hibiscus (+36 more)* |
| 30460-92-5 | Pinocarvone | Terpenoids | NPClassifier, LOTUS, KEGG, LIPID MAPS | Not listed in SPECIATE | 121719 | No | No | Biogenic | Yes | *Brassica; Beta; Laurus; Lachnophyllum; Rubus;* |

| | | | | | | | | | | |
|---|---|---|---|---|---|---|---|---|---|---|
| | | | | | | | | | | *Rumex; Rhynchosia; Lessertia (+58 more)* |
| 11028-42-5 | Alpha-cedrene | Terpenes | NPClassifier, LOTUS | Not listed in SPECIATE | 521207 | No | No | Biogenic | Yes | *Passiflora; Cichorium; Prunus; Rheum; Hippophae; Glycyrrhiza; Actinidia; Vaccinium (+9 more)* |
| 546-28-1 | Beta-cedrene | Terpenes (Terpenoids) | NPClassifier | Not listed in SPECIATE | 11106485 | No | No | Biogenic | Yes | *Brassica; Cinnamomum; Cynara; Mallotus; Mentha; Origanum; Citrus; Dracocephalum (+42 more)* |
| 7785-26-4 | Alpha-pinene | Terpenes | NPClassifier, KEGG | 29 profiles (9 biogenic); top sectors: Biomass Burning; Wildfire (22); Biomass Burning; Prescribed Fire (4); Consumer Products (3) | 6654 | Yes | Yes | Biogenic (dual-source compound) | Yes | *Tamarindus; Brassica; Theobroma; Amelanchier; Vaccinium; Prunus; Oryza; Thymus (+84 more)* |
| 29548-02-5 | Artemisiatriene | Terpenes (Aliphatic hydrocarbons) | NPClassifier | Not listed in SPECIATE | 564723 | No | No | Biogenic | Yes | *Lantana* |
| 555-10-2 | Beta-phellandrene | Terpenes | NPClassifier, LOTUS | 3 profiles; top sectors: Miscellaneous (1); Pulp And Paper; Dryer (1); Pulp And Paper (1) | 11142 | Yes | No | Biogenic | Yes | *Macadamia; Manihot; Durio; Lablab; Helianthus; Vaccinium; Opuntia; Phaseolus (+99 more)* |
| 127-91-3 | Beta-pinene | Terpenes | NPClassifier, KEGG | 122 profiles (9 biogenic); top sectors: Mobile; Onroad; Light Duty (25); Mobile; Onroad (24); Pulp And Paper (22) | 14896 | Yes | Yes | Biogenic (dual-source compound) | Yes | *Durio; Lablab; Helianthus; Byrsonima; Macadamia; Vaccinium; Opuntia; Allium (+74 more)* |
| 99-84-3 | Beta-terpinene | Terpenes | NPClassifier | Not listed in SPECIATE | 66841 | No | No | Biogenic | Yes | *Corylus; Aster; Glycyrrhiza; Sesbania; Sicyos; Brassica;* |

| | | | | | | | | | | |
|---|---|---|---|---|---|---|---|---|---|---|
| | | | | | | | | | | *Aconitum; Artocarpus (+23 more)* |
| 79-92-5 | Camphene | Terpenes | NPClassifier, LOTUS, KEGG | 6 profiles; top sectors: Biomass Burning; Residential Wood Combustion; Woodstove (6) | 6616 | Yes | Yes | Biogenic (dual-source compound) | Yes | *Cymbopogon; Juglans; Morus; Prunus; Ribes; Solanum; Tamarindus; Apium (+86 more)* |
| 18368-95-1 | P-mentha-1,3,8-triene | Terpenes | NPClassifier | Not listed in SPECIATE | 176983 | No | No | Biogenic | Yes | *Cyperus; Silphium; Nephelium; Eruca; Paeonia; Momordica; Myristica; Pastinaca (+23 more)* |
| 3387-41-5 | Sabinene | Terpenes | NPClassifier, KEGG | Not listed in SPECIATE | 18818 | No | No | Biogenic | Yes | *Byrsonima; Rubus; Sambucus; Vaccinium; Vigna; Macadamia; Capsicum; Cyperus (+90 more)* |
| 4501-58-0 | Alpha-campholenal | Terpenoids (Oxygenated aliphatics) | NPClassifier | Not listed in SPECIATE | 1252759 | No | No | Biogenic | Yes | *Chrysanthemum; Solanum; Arctium; Pyrus; Nephelium; Kokoona; Walsura; Ormosia (+4 more)* |
| 586-81-2 | Gamma-terpineol | Terpenoids | NPClassifier | Not listed in SPECIATE | 11467 | No | No | Biogenic | Yes | *Zanthoxylum; Vitis; Helianthus; Teucrium; Ocimum; Phaseolus; Annona; Carum (+36 more)* |
| 24903-95-5 | Nopinone | Terpenoids | NPClassifier, LOTUS | Not listed in SPECIATE | 32735 | No | No | Biogenic | Yes | *Capsicum; Eutrema; Ginkgo; Phaseolus; Zea; Prunus; Brassica; Meiogyne (+4 more)* |
| 513-20-2 | Sabinaketone | Terpenoids (Oxygenated aliphatics) | NPClassifier | Not listed in SPECIATE | 92784 | No | No | Biogenic | Yes | *Vaccinium; Ribes; Mitragyna; Annona;* |

| | | | | | | | | | | |
|---|---|---|---|---|---|---|---|---|---|---|
| | | | | | | | | | | *Convallaria; Zinnia; Melicope; Lafoensia (+8 more)* |
| 1125-12-8 | Thujone | Terpenoids | NPClassifier | Not listed in SPECIATE | 11027 | No | No | Biogenic | Yes | *Haematococcus; Macadamia; Ocimum; Phaseolus; Ornithogalum; Vitex; Vincetoxicum; Ipomoea (+13 more)* |
| 89-83-8 | Thymol | Terpenoids (UNRESOLVED) | NPClassifier, KEGG | Not listed in SPECIATE | 6989 | Yes | Yes | Biogenic (dual-source compound) | Yes | *Pseudotsuga; Viola; Alpinia; Mortonia; Gymnocalycium; Maackia; Betula; Rubia (+86 more)* |
| 3891-98-3 | Farnesane | Terpenes | NPClassifier | 192 profiles; top sectors: Mobile; Onroad (107); Mobile; Onroad; Light Duty (62); Mobile (13) | 19773 | No | No | Biogenic (dual-source compound) | Yes | *Rubus; Punica; Sorghum; Myristica; Mentha; Tetradium* |
| 575-43-9 | Tamarindienal | Terpenoids (Aromatics) | NPClassifier | 13 profiles; top sectors: Mobile (10); Biomass Burning; Residential Wood Combustion; Fireplace (3) | 11328 | Yes | Yes | Biogenic (dual-source compound) | Yes | *Sinapis* |
| 470-82-6 | 1,8-cineole | Terpenoids | NPClassifier, KEGG | 24 profiles (2 biogenic); top sectors: Biomass Burning; Wildfire (18); Consumer Products (5); Composting (1) | 2758 | Yes | Yes | Biogenic (dual-source compound) | Yes | *Oryza; Thymus; Allium; Daucus; Raphanus; Salvia; Manilkara; Angelica (+61 more)* |
| 1795-15-9 | 1-cyclohexyloctane | Aliphatic hydrocarbons | | 139 profiles; top sectors: Mobile; Onroad (77); Mobile; Onroad; Light Duty (50); Mobile; Nonroad (8) | 15712 | No | No | Anthropogenic | Yes | *Ceratonia; Citrus* |

| | | | | | | | | | | |
|---|---|---|---|---|---|---|---|---|---|---|
| 112-41-4 | 1-dodecene | Aliphatic hydrocarbons | | 40 profiles (1 biogenic); top sectors: Mobile; Onroad (19); Biomass Burning; Agriculture (5); Petrochemical; Refinery (5) | 8183 | Yes | Yes | Mixed | Yes | *Vaccinium; Garcinia; Prasinococcus; Sorbus; Vigna; Eragrostis; Benincasa; Ceratonia (+5 more)* |
| 124-11-8 | 1-nonene | Aliphatic hydrocarbons | | 257 profiles (8 biogenic); top sectors: Mobile; Onroad (95); Mobile; Onroad; Light Duty (51); Mobile (51) | 31285 | Yes | Yes | Mixed | Yes | *Pseudocydonia; Gnetum; Psilostrophe; Angelica; Artocarpus; Meum; Ocimum; Ginkgo (+12 more)* |
| 13360-61-7 | 1-pentadecene | Aliphatic hydrocarbons | | Not listed in SPECIATE | 25913 | No | No | Biogenic | Yes | *Trilophozia; Capsicum; Sorbaria; Ricinus; Iris; Garcinia; Polygala; Cicer (+33 more)* |
| 15232-85-6 | 1-pentyl-cyclohexene | Aliphatic hydrocarbons | | Not listed in SPECIATE | 139915 | No | No | Undetermined | – | *nan* |
| 1120-36-1 | 1-tetradecene | Aliphatic hydrocarbons | | 17 profiles; top sectors: Mobile (10); Consumer Products (6); Volatile Chemical Products (1) | 14260 | Yes | Yes | Mixed | Yes | *Grona; Salvia; Aralidium; Psorospermum; Eucalyptus; Ocimum; Psidium; Humulus (+24 more)* |
| 2437-56-1 | 1-tridecene | Aliphatic hydrocarbons | | 15 profiles (1 biogenic); top sectors: Biomass Burning; Agriculture (5); Mobile; Onroad (4); Pulp And Paper; Boiler (3) | 17095 | No | Yes | Mixed | Yes | *Garcinia; Echinacea; Datura; Monodora; Dimocarpus; Primula; Papaver; Chlamydomonas (+3 more)* |
| 2243-98-3 | 1-undecyne | Aliphatic hydrocarbons | | 2 profiles; top sectors: Consumer Products (1); Graphic Arts (1) | 75249 | No | No | Undetermined | Yes | *Cichorium; Rhododendron* |
| 13475-82-6 | 2,2,4,6,6-pentamethylheptane | Aliphatic hydrocarbons | | 25 profiles; top sectors: Consumer | 26058 | No | No | Anthropogenic | Yes | *Grewia; Perezia; Triticum; Dysphania* |

| | | | | | | | | | | |
|---|---|---|---|---|---|---|---|---|---|---|
| | | | | Products (11); Mobile (8); Volatile Chemical Products (4) | | | | | | |
| 192823-15-7 | 2,3,5,8-tetramethyldecane | Aliphatic hydrocarbons | | Not listed in SPECIATE | 545611 | No | No | Undetermined | Yes | *Citrus; Excoecaria; Amomum; Pogostemon; Momordica; Eriobotrya; Hippobroma; Psilotum* |
| 62108-27-4 | 2,4,6-trimethyldecane | Aliphatic hydrocarbons | | Not listed in SPECIATE | 537327 | No | No | Undetermined | – | *nan* |
| 2801-84-5 | 2,4-dimethyldecane | Aliphatic hydrocarbons | | 7 profiles; top sectors: Surface Coating (7) | 520357 | No | No | Anthropogenic | – | *nan* |
| 62108-22-9 | 2,5,9-trimethyldecane | Aliphatic hydrocarbons | | Not listed in SPECIATE | 522020 | No | No | Undetermined | Yes | *Corylus; Pterodon; Rubus* |
| 17302-27-1 | 2,5-dimethylnonane | Aliphatic hydrocarbons | | 1 profiles; top sectors: Surface Coatings (1) | 28456 | No | No | Undetermined | – | *nan* |
| 17301-22-3 | 2,5-dimethylundecane | Aliphatic hydrocarbons | | Not listed in SPECIATE | 28452 | No | No | Undetermined | – | *nan* |
| 31295-56-4 | 2,6,11-trimethyldodecane | Aliphatic hydrocarbons | | Not listed in SPECIATE | 35768 | No | No | Undetermined | Yes | *Syringa; Oryza; Hypericum; Phaseolus; Strychnos* |
| 62108-26-3 | 2,6,8-trimethyldecane | Aliphatic hydrocarbons | | Not listed in SPECIATE | 545608 | No | No | Undetermined | Yes | *Argentina; Smilax* |
| 13150-81-7 | 2,6-dimethyldecane | Aliphatic hydrocarbons | | 9 profiles; top sectors: Surface Coating (8); Surface Coatings (1) | 139395 | No | No | Anthropogenic | – | *nan* |
| 17302-28-2 | 2,6-dimethylnonane | Aliphatic hydrocarbons | | 40 profiles; top sectors: Surface Coating (15); Mobile (10); Volatile Chemical Products (9) | 28457 | No | No | Anthropogenic | Yes | *Garcinia; Maclura* |
| 74645-98-0 | 2,7,10-trimethyldodecane | Aliphatic hydrocarbons | | Not listed in SPECIATE | 93447 | No | No | Undetermined | Yes | *Isodon; Epimedium; Angelica* |
| 17301-25-6 | 2,8-dimethylundecane | Aliphatic hydrocarbons | | Not listed in SPECIATE | 519384 | No | No | Undetermined | – | *nan* |

| | | | | | | | | | | |
|---|---|---|---|---|---|---|---|---|---|---|
| 1560-96-9 | 2-methyltridecane | Aliphatic hydrocarbons | | 86 profiles; top sectors: Consumer Products (66); Volatile Chemical Products (9); Mobile; Aircraft (5) | 15269 | No | No | Anthropogenic | Yes | *Galanthus; Capsicum; Arabidopsis; Citrus; Pinus* |
| 7045-71-8 | 2-methylundecane | Aliphatic hydrocarbons | | 13 profiles; top sectors: Surface Coating (10); Consumer Products (2); Surface Coatings (1) | 23459 | No | No | Anthropogenic | Yes | *Asimina; Brassica; Camellia; Bistorta; Citrullus* |
| 17312-53-7 | 3,6-dimethyldecane | Aliphatic hydrocarbons | | 9 profiles; top sectors: Surface Coating (9) | 519395 | No | No | Anthropogenic | – | *nan* |
| 15869-94-0 | 3,6-dimethyloctane | Aliphatic hydrocarbons | | 428 profiles; top sectors: Mobile (398); Surface Coating (16); Mobile; Onroad (13) | 85927 | No | No | Anthropogenic | – | *nan* |
| 17312-54-8 | 3,7-dimethyldecane | Aliphatic hydrocarbons | | 5 profiles; top sectors: Surface Coating (4); Degreasing (1) | 28468 | No | No | Anthropogenic | No | *nan* |
| 13151-34-3 | 3-methyldecane | Aliphatic hydrocarbons | | 41 profiles; top sectors: Surface Coating (15); Mobile (14); Volatile Chemical Products (7) | 92239 | No | No | Anthropogenic | Yes | *Erysimum; Vaccinium; Eriobotrya* |
| 18435-22-8 | 3-methyltetradecane | Aliphatic hydrocarbons | | Not listed in SPECIATE | 86735 | No | No | Undetermined | Yes | *Citrus; Fragaria; Santalum* |
| 6418-41-3 | 3-methyltridecane | Aliphatic hydrocarbons | | Not listed in SPECIATE | 110848 | No | No | Biogenic | Yes | *Dioscorea; Bistorta; Panax; Tetradium; Millettia; Pimelea; Juniperus; Pterodiscus (+4 more)* |
| 14850-22-7 | 3-octene | Aliphatic hydrocarbons | | Not listed in SPECIATE | 5362722 | No | No | Undetermined | Yes | *Euterpe; Laburnum* |
| 17302-23-7 | 4,5-dimethylnonane | Aliphatic hydrocarbons | | Not listed in SPECIATE | 86541 | No | No | Undetermined | Yes | *Ononis* |

| | | | | | | | | | | |
|---|---|---|---|---|---|---|---|---|---|---|
| 61141-72-8 | 4,6-dimethyldodecane | Aliphatic hydrocarbons | | Not listed in SPECIATE | 545627 | No | No | Undetermined | Yes | *Illicium* |
| 17301-32-5 | 4,7-dimethylundecane | Aliphatic hydrocarbons | | Not listed in SPECIATE | 519389 | No | No | Undetermined | Yes | *Oreocome; Vaccinium; Armoracia; Syzygium; Cichorium; Brassica; Monanthotaxis* |
| 2847-72-5 | 4-methyldecane | Aliphatic hydrocarbons | | 30 profiles; top sectors: Surface Coating (14); Volatile Chemical Products (7); Mobile (4) | 17835 | No | No | Anthropogenic | Yes | *Morus; Artemisia; Ligusticum; Canarium; Putranjiva; Euphorbia; Jurinea* |
| 6117-97-1 | 4-methyldodecane | Aliphatic hydrocarbons | | 8 profiles; top sectors: Surface Coating (7); Degreasing (1) | 521958 | No | No | Anthropogenic | – | *nan* |
| 25117-24-2 | 4-methyltetradecane | Aliphatic hydrocarbons | | Not listed in SPECIATE | 520179 | No | No | Undetermined | Yes | *Zingiber; Triticum; Vaccinium; Zea; Ethulia* |
| 26730-12-1 | 4-methyltridecane | Aliphatic hydrocarbons | | Not listed in SPECIATE | 117325 | No | No | Undetermined | Yes | *Paeonia; Punica; Zingiber; Chrysanthemum; Allium; Cucumis* |
| 2980-69-0 | 4-methylundecane | Aliphatic hydrocarbons | | 16 profiles; top sectors: Surface Coating (11); Mobile (4); Surface Coatings (1) | 520454 | No | No | Anthropogenic | Yes | *Brosimum; Castanea; Bruguiera; Celastrus* |
| 1636-43-7 | 5,6-dimethyldecane | Aliphatic hydrocarbons | | Not listed in SPECIATE | 519255 | No | No | Undetermined | – | *nan* |
| 17312-75-3 | 5-methyl-5-propylnonane | Aliphatic hydrocarbons | | Not listed in SPECIATE | 551397 | No | No | Undetermined | – | *nan* |
| 15869-85-9 | 5-methylnonane | Aliphatic hydrocarbons | | 367 profiles; top sectors: Mobile (355); Mobile; Onroad (8); Surface Coating (4) | 27518 | No | No | Anthropogenic | – | *nan* |
| 41446-61-1 | 6z-tetradecene | Aliphatic hydrocarbons | | Not listed in SPECIATE | 5364500 | No | No | Undetermined | – | *nan* |
| 1678-93-9 | Butylcyclohexane | Aliphatic hydrocarbons | | 130 profiles; top sectors: Mobile (50); Mobile; Onroad (41); | 15506 | No | No | Anthropogenic | Yes | *Vitis; Cydonia; Ipomoea; Triticum; Rheum; Mosla* |

| | | | | | | | | | | |
|---|---|---|---|---|---|---|---|---|---|---|
| | | | | Surface Coating (16) | | | | | | |
| 544-25-2 | Cyclohepta-1,3,5-triene | Aliphatic hydrocarbons | | Not listed in SPECIATE | 11000 | No | No | Undetermined | – | *nan* |
| 124-18-5 | Decane | Aliphatic hydrocarbons | | 1370 profiles (9 biogenic); top sectors: Mobile (549); Mobile; Onroad (216); Oil And Gas; Pond (157) | 15600 | Yes | Yes | Mixed | Yes | *Myrcia; Juglans; Onopordum; Ribes; Artocarpus; Petasites; Eruca; Cicer (+37 more)* |
| 112-40-3 | Dodecane | Aliphatic hydrocarbons | | 1030 profiles (2 biogenic); top sectors: Mobile (435); Mobile; Onroad (193); Mobile; Onroad; Light Duty (133) | 8182 | Yes | Yes | Mixed | Yes | *Veronica; Nicotiana; Daphniphyllum; Asparagus; Juglans; Mentha; Sambucus; Vaccinium (+70 more)* |
| 1678-91-7 | Ethylcyclohexane | Aliphatic hydrocarbons | | 176 profiles; top sectors: Mobile (84); Mobile; Onroad (26); Mobile; Nonroad (14) | 15504 | No | No | Anthropogenic | Yes | *Lupinus; Citrus; Arctium; Glycine* |
| 696-29-7 | Isopropylcyclohexane | Aliphatic hydrocarbons | | 229 profiles; top sectors: Mobile; Onroad (100); Mobile; Onroad; Light Duty (94); Mobile (16) | 12763 | No | No | Anthropogenic | Yes | *Zea* |
| 111-84-2 | Nonane | Aliphatic hydrocarbons | | 1454 profiles (11 biogenic); top sectors: Mobile (561); Mobile; Onroad (233); Oil And Gas; Pond (156) | 8141 | Yes | Yes | Mixed | Yes | *Opuntia; Isodon; Salix; Asparagus; Helianthus; Manihot; Mentha; Origanum (+53 more)* |
| 629-62-9 | Pentadecane | Aliphatic hydrocarbons | | 401 profiles (4 biogenic); top sectors: Mobile; Onroad (184); Consumer Products (66); Mobile; Onroad; Light Duty (61) | 12391 | Yes | Yes | Mixed | Yes | *Salvia; Stachys; Dracocephalum; Lavandula; Boschniakia; Artemisia; Bruguiera; Magnolia (+101 more)* |
| 638-36-8 | Phytane | Aliphatic hydrocarbons (Terpenes) | | 178 profiles; top sectors: Mobile; Onroad (101); Mobile; Onroad; Light Duty (62); | 12523 | Yes | Yes | Mixed | Yes | *Gnetum; Ouratea; Daucus; Eucalyptus; Rubus; Corylus; Rumex; Gypsophila* |

| | | | | | | | | | | |
|---|---|---|---|---|---|---|---|---|---|---|
| | | | | Mobile; Nonroad (8) | | | | | | |
| 2114-42-3 | Prop-2-enylcyclohexane | Aliphatic hydrocarbons | | Not listed in SPECIATE | 75027 | No | No | Undetermined | Yes | *Spinacia* |
| 1678-92-8 | Propyl-cyclohexane | Aliphatic hydrocarbons | | 411 profiles; top sectors: Mobile (378); Surface Coating (17); Mobile; Onroad (11) | 15505 | No | No | Anthropogenic | Yes | *Lathyrus; Hordeum; Mentha; Glycine; Archidendron; Mikania* |
| 629-59-4 | Tetradecane | Aliphatic hydrocarbons | | 369 profiles (4 biogenic); top sectors: Mobile; Onroad (126); Consumer Products (66); Mobile; Onroad; Light Duty (58) | 12389 | Yes | Yes | Mixed | Yes | *Euphorbia; Salvia; Bruguiera; Aegiceras; Cyperus; Petasites; Acorus; Conocephalum (+101 more)* |
| 493-02-7 | Trans-decalin | Aliphatic hydrocarbons (UNRESOLVED) | | 3 profiles; top sectors: Consumer Products (1); Surface Coating (1); Graphic Arts (1) | 7044 | Yes | Yes | Mixed | Yes | *Tanacetum; Fraxinus; Nicotiana; Cucumis* |
| 629-50-5 | Tridecane | Aliphatic hydrocarbons | | 345 profiles (3 biogenic); top sectors: Mobile; Onroad (106); Consumer Products (75); Mobile; Onroad; Light Duty (34) | 12388 | Yes | Yes | Mixed | Yes | *Sempervivum; Manihot; Geum; Ranunculus; Trillium; Salix; Asplenium; Caryocar (+79 more)* |
| 1120-21-4 | Undecane | Aliphatic hydrocarbons | | 1176 profiles (9 biogenic); top sectors: Mobile (493); Mobile; Onroad (155); Mobile; Onroad; Light Duty (115) | 14257 | Yes | Yes | Mixed | Yes | *Baptisia; Mentha; Toona; Oenothera; Marsypianthes; Zinnia; Artemisia; Eucalyptus (+80 more)* |
| 526-73-8 | 1,2,3-trimethylbenzene | Aromatics | | 1101 profiles (13 biogenic); top sectors: Mobile (554); Mobile; Onroad (137); Oil And Gas; Pond (126) | 10686 | Yes | Yes | Mixed | Yes | *Cynomorium; Capsicum; Pinus; Persea; Citrullus; Cucurbita; Cynara; Brassica (+11 more)* |
| 135-77-3 | 1,2,4-trimethoxybenzene | Aromatics | | Not listed in SPECIATE | 67284 | No | No | Undetermined | Yes | *Pinellia* |

| | | | | | | | | | | |
|---|---|---|---|---|---|---|---|---|---|---|
| 95-63-6 | 1,2,4-trimethylbenzene | Aromatics | | 1247 profiles (14 biogenic); top sectors: Mobile (575); Oil And Gas; Pond (154); Mobile; Onroad (150) | 7247 | Yes | Yes | Mixed | Yes | *Juglans; Opuntia; Zizania; Eupatorium; Mallotus; Raphanus; Eugenia; Ficus (+2 more)* |
| 108-67-8 | 1,3,5-trimethylbenzene | Aromatics | | 1335 profiles (15 biogenic); top sectors: Mobile (576); Mobile; Onroad (234); Oil And Gas; Pond (130) | 7947 | Yes | Yes | Mixed | Yes | *Bruguiera; Lathyrus; Cucurbita; Brassica; Persea; Paeonia* |
| 575-37-1 | 1,7-dimethylnaphthalene | Aromatics | | 139 profiles; top sectors: Mobile; Onroad (41); Biomass Burning; Residential Wood Combustion; Fireplace (27); Mobile; Onroad; Light Duty (24) | 11326 | No | No | Anthropogenic | Yes | *Ribes; Prunus* |
| 611-14-3 | 1-ethyl-2-methylbenzene | Aromatics | | 1238 profiles (13 biogenic); top sectors: Mobile (555); Mobile; Onroad (211); Oil And Gas; Pond (119) | 11903 | No | No | Anthropogenic | Yes | *Gymnocarpium; Lonicera; Glycine; Mentha; Vitis; Prunus; Ribes; Oryza (+7 more)* |
| 90-12-0 | 1-methylnaphthalene | Aromatics | | 559 profiles (1 biogenic); top sectors: Mobile; Onroad (220); Mobile; Onroad; Light Duty (125); Consumer Products (53) | 7002 | Yes | Yes | Mixed | Yes | *Medicago; Anthriscus; Vigna; Ficus; Vaccinium; Sorbus; Rubus; Pastinaca (+10 more)* |
| 579-07-7 | 1-phenyl-1,2-propanedione | Aromatics (Oxygenated aliphatics) | | Not listed in SPECIATE | 11363 | No | No | Undetermined | Yes | *Byrsonima; Tillandsia; Rosa; Phyllostachys; Melia; Sorghum; Phytolacca; Benincasa (+1 more)* |
| 26444-19-9 | 2'-methylacetophenone | Aromatics | | Not listed in SPECIATE | 11340 | No | No | Undetermined | Yes | *Vaccinium; Carya; Artocarpus; Pimpinella;* |

| | | | | | | | | | | |
|---|---|---|---|---|---|---|---|---|---|---|
| | | | | | | | | | | *Prunus; Raphanus; Fagopyrum* |
| 96-76-4 | 2,4-di-tert-butylphenol | Aromatics | | Not listed in SPECIATE | 7311 | No | Yes | Mixed | Yes | *Asparagus; Machilus; Phytolacca; Rubus; Ligustrum; Syringa; Hypericum; Lagerstroemia (+11 more)* |
| 24157-81-1 | 2,6-diisopropylnaphthalene | Aromatics | | 30 profiles; top sectors: Consumer Products (30) | 32241 | No | No | Anthropogenic | Yes | *Opuntia* |
| 576-26-1 | 2,6-xylenol | Aromatics | | Not listed in SPECIATE | 11335 | Yes | Yes | Mixed | Yes | *Vigna; Satureja; Allium* |
| 582-24-1 | 2-hydroxyacetophenone | Aromatics (Oxygenated aliphatics) | | Not listed in SPECIATE | 68490 | No | No | Undetermined | Yes | *Corylus; Jodina; Phaseolus* |
| 7786-61-0 | 2-methoxy-4-vinylphenol | Aromatics | | 49 profiles (2 biogenic); top sectors: Biomass Burning; Residential Wood Combustion; Fireplace (22); Biomass Burning; Wildfire (18); Biomass Burning; Residential Wood Combustion; Woodstove (7) | 332 | No | No | Biogenic | Yes | *Mucuna; Sedum; Ilex; Raphanus; Salvia; Juglans; Ambrosia; Gynocardia (+64 more)* |
| 91-57-6 | 2-methylnaphthalene | Aromatics | | 552 profiles (1 biogenic); top sectors: Mobile; Onroad (192); Mobile; Onroad; Light Duty (125); Consumer Products (55) | 7055 | Yes | Yes | Mixed | Yes | *Brassica; Medicago; Phagnalon; Scutellaria; Juglans; Vaccinium; Sicyos; Sinapis (+9 more)* |
| 122-99-6 | 2-phenoxyethanol | Aromatics | | 37 profiles; top sectors: Consumer Products (27); Volatile | 31236 | No | Yes | Anthropogenic | Yes | *Bertholletia; Glechoma; Ficus; Citrus* |

| | | | | | | | | | | |
|---|---|---|---|---|---|---|---|---|---|---|
| | | | | Chemical Products (5); Surface Coating; Architectural (4) | | | | | | |
| 1074-17-5 | 2-propyl toluene | Aromatics | | 743 profiles (7 biogenic); top sectors: Mobile (477); Mobile; Onroad (138); Mobile; Onroad; Light Duty (82) | 14091 | No | No | Anthropogenic | Yes | *Allium; Eutrema; Mentha* |
| 620-14-4 | 3-ethyltoluene | Aromatics | | 1301 profiles (8 biogenic); top sectors: Mobile (572); Mobile; Onroad (229); Oil And Gas; Pond (143) | 12100 | No | No | Anthropogenic | Yes | *Changium; Pediomelum; Rubus; Ajuga; Brassica; Pseudopodospermum; Achillea; Cistus (+1 more)* |
| 13679-41-9 | 3-phenylfuran | Aromatics | | Not listed in SPECIATE | 518802 | No | No | Undetermined | Yes | *Aristolochia; Berberis; Microstrobos; Microula* |
| 104-55-2 | 3-phenylprop-2-enal | Aromatics (Oxygenated aliphatics) | | 28 profiles; top sectors: Biomass Burning; Residential Wood Combustion; Fireplace (21); Biomass Burning; Residential Wood Combustion; Woodstove (7) | 637511 | Yes | Yes | Mixed | Yes | *Carum; Aloe; Hansenia; Conioselinum; Allium; Daucus; Bazzania; Athamanta (+101 more)* |
| 10133-50-3 | 4-isopropenyl benzaldehyde | Aromatics (NOT IN CHEBI) | | Not listed in SPECIATE | 14597914 | No | No | Undetermined | Yes | *Ilex; Artocarpus; Osmorhiza; Betula* |
| 99-89-8 | 4-isopropylphenol | Aromatics | | Not listed in SPECIATE | 7465 | No | No | Undetermined | Yes | *Lupinus; Cynara; Anethum; Apium; Tetragonia* |
| 622-97-9 | 4-methyl styrene | Aromatics | | 35 profiles (7 biogenic); top sectors: Mobile; Onroad (27); Biomass Burning; Prescribed Fire (4); Biomass | 12161 | No | Yes | Anthropogenic | Yes | *Sorghum; Aizoon; Vernonia* |

| | | | | | | | | | | |
|---|---|---|---|---|---|---|---|---|---|---|
| | | | | Burning; Wildfire (4) | | | | | | |
| 5337-93-9 | 4-methylpropiophenone | Aromatics (Oxygenated aliphatics) | | Not listed in SPECIATE | 21429 | No | No | Undetermined | – | *nan* |
| 98-73-7 | 4-t-butylbenzoic acid | Aromatics | | Not listed in SPECIATE | 7403 | No | No | Undetermined | – | *nan* |
| 98-54-4 | 4-tert-butylphenol | Aromatics | | Not listed in SPECIATE | 7393 | No | Yes | Anthropogenic | Yes | *Pouteria* |
| 83-32-9 | Acenaphthene | Aromatics (UNRESOLVED) | | 400 profiles (1 biogenic); top sectors: Mobile; Onroad (170); Mobile; Onroad; Light Duty (105); Biomass Burning; Residential Wood Combustion; Fireplace (33) | 6734 | Yes | Yes | Mixed | Yes | *Cymbopogon* |
| 98-86-2 | Acetophenone | Aromatics (Oxygenated aliphatics) | | 44 profiles (3 biogenic); top sectors: Pulp And Paper (9); Mobile; Onroad (8); Biomass Burning; Residential Wood Combustion; Fireplace (6) | 7410 | Yes | Yes | Mixed | Yes | *Macadamia; Saururus; Ipomoea; Garcinia; Allium; Raphanus; Brassica; Vaccinium (+45 more)* |
| 583-04-0 | Allyl benzoate | Aromatics | | Not listed in SPECIATE | 11406 | No | No | Undetermined | Yes | *Momordica; Carica; Orthosiphon; Avena* |
| 100-52-7 | Benzaldehyde | Aromatics | | 353 profiles (15 biogenic); top sectors: Mobile; Onroad (128); Mobile; Onroad; Light Duty (78); Mobile; Nonroad (27) | 240 | Yes | Yes | Mixed | Yes | *Ribes; Daucus; Brassica; Theobroma; Amelanchier; Aspidosperma; Abies; Rhododendron (+68 more)* |
| 65-85-0 | Benzoic acid | Aromatics | | 116 profiles; top sectors: Mobile; Onroad (42); Mobile; Onroad; Light Duty (23); Biomass Burning; | 243 | Yes | Yes | Mixed | Yes | *Brassica; Cucumis; Cymbopogon; Juglans; Morus; Prunus; Ribes; Solanum (+65 more)* |

| | | | | | | | | | | |
|---|---|---|---|---|---|---|---|---|---|---|
| | | | | Residential Wood Combustion; Fireplace (16) | | | | | | |
| 119-61-9 | Benzophenone | Aromatics (Oxygenated aliphatics) | | Not listed in SPECIATE | 3102 | Yes | Yes | Mixed | Yes | *Amelanchier; Taxus; Capsicum; Genista; Sorbus* |
| 92-52-4 | Biphenyl | Aromatics | | 368 profiles; top sectors: Mobile; Onroad (142); Mobile; Onroad; Light Duty (124); Biomass Burning; Residential Wood Combustion; Fireplace (30) | 7095 | Yes | Yes | Mixed | Yes | *Maackia; Aconitum; Ziziphus; Borago; Vitis; Medinilla; Tragopogon; Paris (+4 more)* |
| 104-51-8 | Butylbenzene | Aromatics | | 279 profiles (7 biogenic); top sectors: Mobile; Onroad (68); Mobile (62); Consumer Products (57) | 7705 | No | Yes | Anthropogenic | Yes | *Solanum; Pimpinella; Chenopodium* |
| 98-82-8 | Cumene | Aromatics | | 1146 profiles (13 biogenic); top sectors: Mobile (525); Mobile; Onroad (159); Oil And Gas; Pond (100) | 7406 | Yes | Yes | Mixed | Yes | *Vaccinium; Vitis; Pistacia; Paeonia; Picrasma; Rhododendron; Akebia; Coffea (+4 more)* |
| 122-03-2 | Cuminaldehyde | Aromatics | | Not listed in SPECIATE | 326 | No | No | Biogenic | Yes | *Oryza; Thymus; Allium; Daucus; Centipeda; Macaranga; Mallotus; Alnus (+78 more)* |
| 132-64-9 | Dibenzofuran | Aromatics | | 325 profiles; top sectors: Mobile; Onroad (164); Mobile; Onroad; Light Duty (100); Biomass Burning; Residential Wood Combustion; Fireplace (16) | 568 | No | Yes | Anthropogenic | Yes | *Ficus; Allium; Sorbus; Phaseolus; Murraya; Helicteres* |

| | | | | | | | | | | |
|---|---|---|---|---|---|---|---|---|---|---|
| 15764-16-6 | Dimethyl benzaldehyde | Aromatics (Oxygenated aliphatics) | | 4 profiles; top sectors: Mobile; Onroad (4) | 61814 | No | No | Anthropogenic | Yes | *Capsicum; Annona; Artemisia; Angelica; Cichorium* |
| 131-11-3 | Dimethyl phthalate | Aromatics | | 18 profiles; top sectors: Consumer Products (5); Petrochemical; Refinery (3); Volatile Chemical Products (2) | 8554 | Yes | Yes | Mixed | Yes | *Cucurbita; Sicyos; Salvia; Allamanda; Sambucus; Forsythia; Actinidia; Trigonella (+4 more)* |
| 27831-13-6 | Dimethylstyrene | Aromatics | | 5 profiles; top sectors: Industrial (5) | 33937 | No | No | Anthropogenic | Yes | *Eucalyptus* |
| 95-93-2 | Durene | Aromatics | | 594 profiles; top sectors: Mobile (449); Mobile; Onroad (51); Mobile; Onroad; Light Duty (49) | 7269 | No | No | Anthropogenic | Yes | *Hippophae; Capsicum* |
| 93-89-0 | Ethyl benzoate | Aromatics | | 2 profiles; top sectors: Agriculture; Silage (2) | 7165 | No | No | Biogenic | Yes | *Morus; Prunus; Byrsonima; Salix; Juglans; Mentha; Durio; Lablab (+40 more)* |
| 100-41-4 | Ethylbenzene | Aromatics | | 1699 profiles (22 biogenic); top sectors: Mobile (588); Mobile; Onroad (251); Oil And Gas; Pond (141) | 7500 | Yes | Yes | Mixed | Yes | *Piper; Lippia; Ulex; Juniperus; Hornschuchia; Asparagus; Juglans; Sorghum (+32 more)* |
| 86-73-7 | Fluorene | Aromatics | | 558 profiles (1 biogenic); top sectors: Mobile; Onroad (277); Mobile; Onroad; Light Duty (125); Biomass Burning; Residential Wood Combustion; Woodstove (45) | 6853 | Yes | Yes | Mixed | Yes | *Asparagus; Diospyros; Passiflora; Aronia* |
| 6789-88-4 | Hexyl benzoate | Aromatics | | Not listed in SPECIATE | 23235 | No | No | Biogenic | Yes | *Crotalaria; Citrus; Bupleurum; Ononis;* |

| | | | | | | | | | | |
|---|---|---|---|---|---|---|---|---|---|---|
| | | | | | | | | | | *Serenoa; Annona; Salvia; Pseudocydonia (+17 more)* |
| 496-11-7 | Indane | Aromatics (UNRESOLVED) | | 825 profiles (9 biogenic); top sectors: Mobile (525); Mobile; Onroad (89); Mobile; Onroad; Light Duty (85) | 10326 | No | No | Anthropogenic | Yes | *Oenothera; Illicium; Matricaria* |
| 4247-02-3 | Isobutylparaben | Aromatics | | Not listed in SPECIATE | 20240 | No | No | Undetermined | Yes | *Phytolacca* |
| 93-16-3 | Isomethyleugenol | Aromatics | | Not listed in SPECIATE | 637776 | No | No | Biogenic | Yes | *Melissa; Trigonella; Citrus; Magnolia; Artemisia; Rheum; Mespilus; Vigna (+31 more)* |
| 585-74-0 | M-methylacetophenone | Aromatics (Oxygenated aliphatics) | | Not listed in SPECIATE | 11455 | No | No | Undetermined | Yes | *Urolepis; Litchi; Salvia; Oenothera; Lappula; Origanum* |
| 620-23-5 | M-tolualdehyde | Aromatics | | 55 profiles (3 biogenic); top sectors: Biomass Burning; Wildfire (19); Mobile; Onroad (15); Biomass Burning; Agriculture (6) | 12105 | No | Yes | Mixed | Yes | *Lablab; Vaccinium; Beta; Tetrastigma; Agave; Brassica; Croton; Carthamus (+2 more)* |
| 104-20-1 | Methoxyphenyl butanone | Aromatics | | Not listed in SPECIATE | 61007 | No | No | Undetermined | Yes | *Adiantum; Cucumis; Pittocaulon; Pimpinella; Cornus; Psidium; Persicaria* |
| 93-58-3 | Methyl benzoate | Aromatics | | 13 profiles; top sectors: Biomass Burning; Residential Wood Combustion; Woodstove (7); Biomass Burning; Residential | 7150 | Yes | Yes | Mixed | Yes | *Xanthosoma; Brassica; Hyacinthoides; Manihot; Macadamia; Durio; Lablab; Helianthus (+97 more)* |

| | | | | | | | | | | |
|---|---|---|---|---|---|---|---|---|---|---|
| | | | | Wood Combustion; Fireplace (4); Agriculture; Silage (2) | | | | | | |
| 119-36-8 | Methyl salicylate | Aromatics | | 8 profiles; top sectors: Consumer Products (6); Biomass Burning; Agriculture (2) | 4133 | Yes | Yes | Mixed | Yes | *Rubia; Picea; Lablab; Cyperus; Artemisia; Gypsophila; Rhodiola; Frullania (+80 more)* |
| 91-20-3 | Naphthalene | Aromatics | | 1286 profiles (12 biogenic); top sectors: Mobile (448); Mobile; Onroad (274); Mobile; Onroad; Light Duty (205) | 931 | Yes | Yes | Mixed | Yes | *Cymbopogon; Amelanchier; Tamarindus; Apium; Vaccinium; Prunus; Oryza; Thymus (+67 more)* |
| 1450-72-2 | Ortho-acetyl-para-cresol | Aromatics (Oxygenated aliphatics) | | Not listed in SPECIATE | 15068 | No | No | Undetermined | Yes | *Laurus; Litchi; Abelmoschus; Momordica; Rubus* |
| 123-11-5 | P-methoxybenzaldehyde | Aromatics | | Not listed in SPECIATE | 31244 | Yes | Yes | Mixed | Yes | *Celmisia; Solanum; Richetia; Gymnema; Psilostrophe; Dracocephalum; Annona; Capsicum (+89 more)* |
| 80-46-6 | P-tert-amylphenol | Aromatics | | Not listed in SPECIATE | 6643 | No | Yes | Anthropogenic | Yes | *Theobroma; Paeonia; Meconopsis; Rauvolfia; Canarium* |
| 104-87-0 | P-tolualdehyde | Aromatics | | 41 profiles (1 biogenic); top sectors: Mobile; Onroad; Light Duty (9); Consumer Products (7); Biomass Burning; Agriculture (5) | 7725 | Yes | Yes | Mixed | Yes | *Sambucus; Emblica; Callicarpa; Panicum; Satureja; Nierembergia; Solanum; Vaccinium (+18 more)* |
| 108-95-2 | Phenol | Aromatics | | 157 profiles (14 biogenic); top sectors: Mobile; | 996 | Yes | Yes | Mixed | Yes | *Prunus; Ribes; Solanum; Tamarindus; Apium;* |

| | | | | | | | | | | |
|---|---|---|---|---|---|---|---|---|---|---|
| | | | | Onroad (29); Biomass Burning; Wildfire (22); Pulp And Paper (17) | | | | | | *Cucumis; Juglans; Eleocharis (+87 more)* |
| 122-78-1 | Phenylacetaldehyde | Aromatics (Oxygenated aliphatics) | | Not listed in SPECIATE | 998 | No | No | Biogenic | Yes | *Daucus; Raphanus; Salvia; Juglans; Eleocharis; Manilkara; Xanthosoma; Brassica (+83 more)* |
| 1074-12-0 | Phenylglyoxal | Aromatics (Oxygenated aliphatics) | | Not listed in SPECIATE | 14090 | No | No | Undetermined | Yes | *Camellia; Myristica; Chrysosplenium; Descurainia; Actaea* |
| 611-73-4 | Phenylglyoxylic acid | Aromatics (Oxygenated aliphatics) | | Not listed in SPECIATE | 11915 | No | No | Undetermined | Yes | *Triticum; Vaccinium; Cicer; Citrus* |
| 85-44-9 | Phthalic anhydride | Aromatics (Oxygenated aliphatics) | | 15 profiles; top sectors: Chemical Manufacturing (4); Petrochemical; Refinery (4); Consumer Products (3) | 6811 | No | Yes | Mixed | Yes | *Alcea; Lablab; Helianthus; Asparagus; Carica; Vaccinium; Citrus; Phytolacca (+3 more)* |
| 103-65-1 | Propylbenzene | Aromatics | | 1313 profiles (14 biogenic); top sectors: Mobile (569); Mobile; Onroad (235); Mobile; Onroad; Light Duty (118) | 7668 | Yes | Yes | Mixed | Yes | *Asparagus; Juglans; Mentha; Asphodelus; Piper; Brassica; Solanum; Glycyrrhiza (+1 more)* |
| 90-02-8 | Salicylaldehyde | Aromatics | | Not listed in SPECIATE | 6998 | Yes | Yes | Mixed | Yes | *Vaccinium; Panax; Brassica; Bryum; Cedrus; Ficus; Basella; Genista (+8 more)* |
| 135-98-8 | Sec-butylbenzene | Aromatics | | 600 profiles; top sectors: Mobile (357); Mobile; Onroad; Light Duty (82); Mobile; Onroad (63) | 8680 | Yes | Yes | Mixed | – | *nan* |

| | | | | | | | | | | |
|---|---|---|---|---|---|---|---|---|---|---|
| 100-42-5 | Styrene | Aromatics | | 472 profiles (16 biogenic); top sectors: Mobile; Onroad (116); Mobile (51); Oil And Gas; Pond (42) | 7501 | Yes | Yes | Mixed | Yes | *Artemisia; Ocimum; Mentha; Byrsonima; Pectinopitys; Meliosma; Picea; Cucurbita (+39 more)* |
| 774-65-2 | Tert-butyl benzoate | Aromatics | | Not listed in SPECIATE | 69886 | No | No | Undetermined | – | *nan* |
| 108-88-3 | Toluene | Aromatics | | 1697 profiles (25 biogenic); top sectors: Mobile (586); Oil And Gas; Pond (165); Mobile; Onroad (163) | 1140 | Yes | Yes | Mixed | Yes | *Ardisia; Bryonia; Mollinedia; Citrus; Physalis; Pangium; Pedicularis; Wikstroemia (+40 more)* |
| 121-33-5 | Vanillin | Aromatics | | 189 profiles; top sectors: Mobile; Onroad (71); Mobile; Onroad; Light Duty (50); Biomass Burning; Residential Wood Combustion; Fireplace (30) | 1183 | Yes | Yes | Mixed | Yes | *Eleocharis; Manilkara; Xanthosoma; Brassica; Lepidium; Theobroma; Amelanchier; Salvia (+41 more)* |
| 769-78-8 | Vinyl benzoate | Aromatics | | Not listed in SPECIATE | 13037 | No | No | Undetermined | Yes | *Morus; Verbena* |
| 108-38-3 | Xylene | Aromatics | | 266 profiles (5 biogenic); top sectors: Mobile (83); Consumer Products (27); Mobile; Onroad (22) | 7929 | Yes | Yes | Mixed | Yes | *Lablab; Helianthus; Vaccinium; Cocos; Nicotiana; Micromeria; Psidium; Punica (+18 more)* |
| 288-39-1 | 1,2,5-thiadiazole | Others (N, S, Halo) (UNRESOLVED) | | Not listed in SPECIATE | 559537 | No | No | Undetermined | – | *nan* |
| 78-87-5 | 1,2-dichloropropane | Others (N, S, Halo) (UNRESOLVED) | | 4 profiles; top sectors: Landfill (1); Pulp And Paper; Boiler (1); Open Burning (1) | 6564 | No | Yes | Anthropogenic | – | *nan* |
| 505-20-4 | 1,2-dithiane | Others (N, S, Halo) | | Not listed in SPECIATE | 136335 | No | No | Undetermined | – | *nan* |
| 616-47-7 | 1-methyl-1h-imidazole | Others (N, S, Halo) (Aromatics) | | Not listed in SPECIATE | 1390 | No | No | Undetermined | – | *nan* |

| | | | | | | | | | | |
|---|---|---|---|---|---|---|---|---|---|---|
| 20348-18-9 | 2-amino-4-methyl-3-pyridinol | Others (N, S, Halo) (Aromatics) | | Not listed in SPECIATE | 580054 | No | No | Undetermined | – | *nan* |
| 18936-17-9 | 2-methylbutanenitrile | Others (N, S, Halo) | | Not listed in SPECIATE | 29339 | No | No | Undetermined | Yes | *Rzedowskia; Psidium; Platycladus* |
| 22990-77-8 | 2-piperidinemethanamine | Others (N, S, Halo) | | Not listed in SPECIATE | 90865 | No | No | Undetermined | – | *nan* |
| 5961-33-1 | 3-methyl-3-phenylazetidine | Others (N, S, Halo) (Aromatics) | | Not listed in SPECIATE | 22249 | No | No | Undetermined | Yes | *Theobroma* |
| 767-00-0 | 4-cyanophenol | Others (N, S, Halo) (Aromatics) | | Not listed in SPECIATE | 13019 | No | No | Undetermined | – | *nan* |
| 100-47-0 | Benzonitrile | Others (N, S, Halo) (Aromatics) | | 26 profiles (9 biogenic); top sectors: Biomass Burning; Wildfire (22); Biomass Burning; Prescribed Fire (4) | 7505 | No | Yes | Mixed | Yes | *Manihot* |
| 611-20-1 | Benzonitrile, 2-hydroxy- | Others (N, S, Halo) (Aromatics) | | Not listed in SPECIATE | 11907 | No | No | Undetermined | – | *nan* |
| 95-16-9 | Benzothiazole | Others (N, S, Halo) | | 4 profiles; top sectors: Dry Cleaning (2); Consumer Products (1); Graphic Arts (1) | 7222 | No | Yes | Mixed | Yes | *Helianthus; Vaccinium; Byrsonima; Piper; Asparagus; Orthosiphon; Breynia; Erica (+14 more)* |
| 98-88-4 | Benzoyl chloride | Others (N, S, Halo) (Aromatics) | | Not listed in SPECIATE | 7412 | No | Yes | Anthropogenic | Yes | *Illicium* |
| 3012-37-1 | Benzyl thiocyanate | Others (N, S, Halo) | | Not listed in SPECIATE | 18170 | No | No | Undetermined | Yes | *Capsicum; Ziziphus; Vaccinium; Phyllostachys; Populus; Vitis* |
| 79-11-8 | Chloroacetic acid | Others (N, S, Halo) | | Not listed in SPECIATE | 300 | No | Yes | Anthropogenic | Yes | *Prunus; Elsholtzia* |
| 108-90-7 | Chlorobenzene | Others (N, S, Halo) (Aromatics) | | 71 profiles (3 biogenic); top sectors: Pulp And Paper (19); Consumer Products (9); Chemical Manufacturing (8) | 7964 | Yes | Yes | Mixed | Yes | *Solanum* |

| | | | | | | | | | | |
|---|---|---|---|---|---|---|---|---|---|---|
| 38917-61-2 | Dimethyl cyclopentapyrazine | Others (N, S, Halo) | | Not listed in SPECIATE | 530404 | No | No | Undetermined | Yes | *Veratrum; Citrullus; Citrus* |
| 1467-79-4 | Dimethyl-cyanamide | Others (N, S, Halo) | | Not listed in SPECIATE | 15112 | No | No | Undetermined | – | *nan* |
| 645-05-6 | Hexamethylmelamine | Others (N, S, Halo) (Aromatics) | | Not listed in SPECIATE | 2123 | No | No | Undetermined | Yes | *Phaseolus* |
| 6919-61-5 | N-methoxy-n-methylbenzamide | Others (N, S, Halo) | | Not listed in SPECIATE | 569575 | No | No | Undetermined | – | *nan* |
| 1122-85-6 | Phenyl cyanate | Others (N, S, Halo) | | Not listed in SPECIATE | 70740 | No | No | Undetermined | – | *nan* |
| 110-89-4 | Piperidine | Others (N, S, Halo) | | Not listed in SPECIATE | 8082 | Yes | Yes | Mixed | Yes | *Coleus; Piper; Brassica; Vaccinium; Zizania; Crateva; Pimenta; Syzygium (+13 more)* |
| 500-22-1 | Pyridine-3-carbaldehyde | Others (N, S, Halo) | | Not listed in SPECIATE | 10371 | No | No | Undetermined | Yes | *Cucumis; Crateva;* |
| 67-71-0 | Sulfonyldimethane | Others (N, S, Halo) | | 6 profiles; top sectors: Agriculture; Animal (6) | 6213 | No | No | Biogenic | Yes | *Manihot; Vigna; Meum; Salvia; Neslia; Sarracenia; Fragaria; Ribes (+10 more)* |
| 127-18-4 | Tetrachloroethene | Others (N, S, Halo) | | 135 profiles (4 biogenic); top sectors: Consumer Products (60); Pulp And Paper (15); Mobile; Onroad (10) | 31373 | No | Yes | Anthropogenic | Yes | *Prunus; Forsythia* |
| 288-47-1 | Thiazole | Others (N, S, Halo) (Aromatics) | | Not listed in SPECIATE | 9256 | No | No | Undetermined | Yes | *Strychnos; Emblica* |
| 93-55-0 | Triphenylsulfonium | Others (N, S, Halo) (Oxygenated aliphatics) | | Not listed in SPECIATE | 7148 | No | Yes | Anthropogenic | Yes | *Daucus; Manilkara; Melilotus; Gaylussacia; Brassica; Annona* |
| 5077-67-8 | 1-hydroxybutan-2-one | Oxygenated aliphatics | | Not listed in SPECIATE | 521300 | No | No | Undetermined | Yes | *Ephedra; Brassica; Cynoglossum; Mentha; Erythrina; Pistacia; Euphorbia; Syncolostemon* |

| | | | | | | | | | | |
|---|---|---|---|---|---|---|---|---|---|---|
| 4312-99-6 | 1-octen-3-one | Oxygenated aliphatics (UNRESOLVED) | | Not listed in SPECIATE | 61346 | No | No | Biogenic | Yes | *Marsdenia; Eugenia; Canarium; Dendrobium; Cichorium; Hippophae; Curcuma; Pulicaria (+13 more)* |
| 15045-43-9 | 2,2,5,5-tetramethyltetrahydrofuran | Oxygenated aliphatics | | Not listed in SPECIATE | 27010 | No | No | Undetermined | Yes | *Capsicum* |
| 6570-88-3 | 2,3,4-trimethyl-1-pentanol | Oxygenated aliphatics (UNRESOLVED) | | Not listed in SPECIATE | 550859 | No | No | Undetermined | – | *nan* |
| 2816-57-1 | 2,6-dimethylcyclohexanone | Oxygenated aliphatics (UNRESOLVED) | | Not listed in SPECIATE | 17780 | No | No | Undetermined | Yes | *Iberis; Carica* |
| 16400-72-9 | 2-dodecen-5-olide | Oxygenated aliphatics | | Not listed in SPECIATE | 61834 | No | No | Undetermined | Yes | *Allium* |
| 104-76-7 | 2-ethylhexan-1-ol | Oxygenated aliphatics (UNRESOLVED) | | 27 profiles; top sectors: Consumer Products (13); Volatile Chemical Products (4); Agriculture; Animal (3) | 7720 | Yes | Yes | Mixed | Yes | *Cinnamodendron; Lysimachia; Citrullus; Citrus; Cydonia; Ipomoea; Levisticum; Tagetes (+21 more)* |
| 123-05-7 | 2-ethylhexanal | Oxygenated aliphatics | | Not listed in SPECIATE | 31241 | Yes | Yes | Mixed | Yes | *Nicotiana; Tacca; Cichorium* |
| 149-57-5 | 2-ethylhexanoic acid | Oxygenated aliphatics | | Not listed in SPECIATE | 8697 | No | Yes | Anthropogenic | Yes | *Cucurbita* |
| 505-57-7 | 2-hexenal | Oxygenated aliphatics | | Not listed in SPECIATE | 5281168 | No | No | Biogenic | Yes | *Annona; Carum; Daucus; Erythrina; Ferula; Gossypium; Vitis; Nephelium (+98 more)* |
| 54845-28-2 | 2-hexenyl 2-hexenoate | Oxygenated aliphatics (NOT IN CHEBI) | | Not listed in SPECIATE | 5369062 | No | No | Undetermined | – | *nan* |
| 2445-77-4 | 2-methylbutyl isovalerate | Oxygenated aliphatics | | Not listed in SPECIATE | 62445 | No | No | Undetermined | Yes | *Abelmoschus; Annona; Carum; Vigna; Myrtus* |
| 3777-69-3 | 2-pentylfuran | Oxygenated aliphatics | | 2 profiles; top sectors: Agriculture; Silage (2) | 19602 | No | No | Biogenic | Yes | *Capsicum; Ziziphus; Basella; Secale; Swertia;* |

| | | | | | | | | | | |
|---|---|---|---|---|---|---|---|---|---|---|
| | | | | | | | | | | *Zanthoxylum; Solanum; Skimmia (+73 more)* |
| 53448-07-0 | 2-undecenal | Oxygenated aliphatics | | 7 profiles (4 biogenic); top sectors: Cooking; Frying (5); Cooking; Charbroiling (2) | 5283356 | No | No | Biogenic | Yes | *Triticum; Dimocarpus; Allium; Senna; Leibnitzia; Tarenna; Pteridium; Vigna (+7 more)* |
| 1123-19-9 | 3-acetyl-3-methyldihydrofuran-2(3h)-one | Oxygenated aliphatics | | Not listed in SPECIATE | 79144 | No | No | Undetermined | – | *nan* |
| 498-60-2 | 3-furaldehyde | Oxygenated aliphatics | | 12 profiles (7 biogenic); top sectors: Biomass Burning; Prescribed Fire (4); Biomass Burning; Wildfire (4); Biomass Burning; Residential Wood Combustion; Fireplace (3) | 10351 | No | No | Biogenic | Yes | *Thespesia; Cynara; Thymus* |
| 1193-18-6 | 3-methyl-2-cyclohexen-1-one | Oxygenated aliphatics (Aliphatic hydrocarbons) | | Not listed in SPECIATE | 14511 | No | No | Undetermined | Yes | *Angylocalyx; Ceratiola* |
| 591-24-2 | 3-methylcyclohexanone | Oxygenated aliphatics | | Not listed in SPECIATE | 11567 | No | No | Undetermined | Yes | *Rehmannia; Veronica; Secale; Brassica; Ananas; Ophrys; Actinidia; Allium (+1 more)* |
| 1757-42-2 | 3-methylcyclopentanone | Oxygenated aliphatics | | Not listed in SPECIATE | 15650 | No | No | Undetermined | Yes | *Triticum; Rubus* |
| 589-92-4 | 4-methylcyclohexanone | Oxygenated aliphatics | | Not listed in SPECIATE | 11525 | No | No | Undetermined | Yes | *Tephroseris; Daucus; Isodon; Eucommia; Allium* |
| 2492-43-5 | 4-oxohex-2-enal | Oxygenated aliphatics | | Not listed in SPECIATE | 6365145 | No | No | Undetermined | Yes | *Cichorium; Epimedium* |
| 20019-64-1 | 5,5-dimethyl-2(5h)-furanone | Oxygenated aliphatics | | Not listed in SPECIATE | 29909 | No | No | Undetermined | Yes | *Cistus; Callitropsis* |

| | | | | | | | | | | |
|---|---|---|---|---|---|---|---|---|---|---|
| 31502-19-9 | 6e-nonen-1-ol | Oxygenated aliphatics (UNRESOLVED) | | Not listed in SPECIATE | 5362811 | No | No | Undetermined | Yes | *Cirsium* |
| 626-82-4 | Butyl hexanoate | Oxygenated aliphatics | | Not listed in SPECIATE | 12294 | No | No | Biogenic | Yes | *Hypericum; Xylopia; Agave; Triticum; Pinus; Mentha; Artemisia; Mandragora (+4 more)* |
| 500-02-7 | Crypton | Oxygenated aliphatics (NOT IN CHEBI) | | Not listed in SPECIATE | 92780 | No | No | Biogenic | Yes | *Ocimum; Phaseolus; Lathyrus; Arabidopsis; Cirsium; Limnophila; Corchorus; Allium (+25 more)* |
| 108-94-1 | Cyclohexanone | Oxygenated aliphatics | | 86 profiles (3 biogenic); top sectors: Consumer Products (29); Biomass Burning; Wildfire (18); Agriculture; Silage (7) | 7967 | No | Yes | Mixed | Yes | *Garcinia; Clusia; Brassica; Juniperus; Carthamus; Vaccinium; Abelmoschus; Annona (+15 more)* |
| 120-92-3 | Cyclopentanone | Oxygenated aliphatics | | 30 profiles (9 biogenic); top sectors: Biomass Burning; Wildfire (22); Biomass Burning; Prescribed Fire (4); Biomass Burning; Residential Wood Combustion; Fireplace (3) | 8452 | No | Yes | Mixed | Yes | *Avena; Paeonia; Phellodendron; Citrus; Cuminum; Albizia; Nephelium; Eruca (+94 more)* |
| 3913-81-3 | Dec-2-enal | Oxygenated aliphatics | | Not listed in SPECIATE | 5283345 | No | No | Biogenic | Yes | *Coreopsis; Juglans; Vaccinium; Brassica; Sicyos; Arctium; Erycibe; Trifolium (+43 more)* |
| 112-31-2 | Decanal | Oxygenated aliphatics | | 45 profiles (4 biogenic); top sectors: Biomass Burning; Wildfire (18); | 8175 | Yes | Yes | Mixed | Yes | *Asparagus; Juglans; Mentha; Rubus; Sambucus; Vigna;* |

| | | | | | | | | | | |
|---|---|---|---|---|---|---|---|---|---|---|
| | | | | Agriculture; Animal (5); Mobile; Aircraft (4) | | | | | | *Macadamia; Manihot (+89 more)* |
| 334-48-5 | Decanoic acid | Oxygenated aliphatics | | 175 profiles (11 biogenic); top sectors: Mobile; Onroad (66); Mobile; Onroad; Light Duty (35); Biomass Burning; Residential Wood Combustion; Woodstove (21) | 2969 | Yes | Yes | Mixed | Yes | *Cymbopogon; Juglans; Morus; Prunus; Ribes; Brassica; Theobroma; Amelanchier (+74 more)* |
| 112-34-5 | Diethylene glycol monobutyl ether | Oxygenated aliphatics | | 113 profiles; top sectors: Consumer Products (58); Surface Coating; Architectural (9); Degreasing (6) | 8177 | No | Yes | Anthropogenic | Yes | *Vaccinium* |
| 112-54-9 | Dodecanal | Oxygenated aliphatics | | 7 profiles; top sectors: Mobile; Onroad (3); Cooking; Charbroiling (2); Mobile; Nonroad (1) | 8194 | No | No | Mixed | Yes | *Anisomeles; Gouania; Allamanda; Helianthus; Tamilnadia; Artemisia; Euphorbia; Cinnamomum (+50 more)* |
| 143-07-7 | Dodecanoic acid | Oxygenated aliphatics | | 194 profiles (12 biogenic); top sectors: Mobile; Onroad (55); Mobile; Onroad; Light Duty (48); Biomass Burning; Residential Wood Combustion; Fireplace (32) | 3893 | Yes | Yes | Mixed | Yes | *Daucus; Raphanus; Salvia; Juglans; Eleocharis; Manilkara; Xanthosoma; Mentha (+11 more)* |
| 105-60-2 | Epsilon-caprolactam | Oxygenated aliphatics (Others (N, S, Halo)) | | Not listed in SPECIATE | 7768 | Yes | Yes | Mixed | Yes | *Annona; Dysphania; Arctium; Carica; Cichorium; Canarium; Ulva; Basella* |
| 2983-37-1 | Ethyl 2-ethylhexanoate | Oxygenated aliphatics | | Not listed in SPECIATE | 102916 | No | No | Undetermined | Yes | *Verbena* |

| | | | | | | | | | | |
|---|---|---|---|---|---|---|---|---|---|---|
| 695-06-7 | Gamma-caprolactone | Oxygenated aliphatics | | 5 profiles (2 biogenic); top sectors: Cooking; Frying (3); Cooking; Charbroiling (2) | 12756 | No | No | Biogenic | Yes | *Helichrysum; Aglaia; Parastrephia; Daphniphyllum; Porella; Salix; Rehmannia; Camellia (+8 more)* |
| 110-43-0 | Heptan-2-one | Oxygenated aliphatics | | 54 profiles; top sectors: Consumer Products (14); Volatile Chemical Products (9); Surface Coating; Architectural (6) | 8051 | Yes | Yes | Mixed | Yes | *Morus; Lippia; Asparagus; Juglans; Mentha; Rubus; Sambucus; Vaccinium (+52 more)* |
| 106-35-4 | Heptan-3-one | Oxygenated aliphatics | | 1 profiles; top sectors: Agriculture; Animal (1) | 7802 | Yes | Yes | Mixed | Yes | *Eugenia; Tinospora; Vaccinium; Vigna; Camellia; Cocos* |
| 111-71-7 | Heptanal | Oxygenated aliphatics | | 48 profiles (7 biogenic); top sectors: Biomass Burning; Wildfire (18); Mobile; Onroad (9); Agriculture; Animal (6) | 8130 | Yes | Yes | Mixed | Yes | *Garcinia; Artemisia; Peripterygia; Syzygiella; Taraxacum; Centaurea; Diospyros; Tabernaemontana (+59 more)* |
| 111-14-8 | Heptanoic acid | Oxygenated aliphatics | | 58 profiles (3 biogenic); top sectors: Mobile; Onroad; Light Duty (28); Mobile; Onroad (16); Agriculture; Animal (6) | 8094 | Yes | Yes | Mixed | Yes | *Eucalyptus; Stauntonia; Diospyros; Arbutus; Stemodia; Bowdichia; Viguiera; Baccharis (+98 more)* |
| 78-59-1 | Isophorone | Oxygenated aliphatics | | 7 profiles; top sectors: Consumer Products (4); Pulp And Paper (2); Surface Coatings; Aerosol (1) | 6544 | Yes | Yes | Mixed | Yes | *Asparagus; Phytolacca; Ilex; Aristolochia; Juniperus; Capsicum; Kandelia; Cucurbita (+15 more)* |
| 57576-09-7 | Isopulegol acetate | Oxygenated aliphatics (Terpenoids) | | Not listed in SPECIATE | 94579 | No | No | Undetermined | Yes | *Crotalaria; Medicago; Balanites; Artemisia;* |

| | | | | | | | | | | |
|---|---|---|---|---|---|---|---|---|---|---|
| | | | | | | | | | | *Adenanthera; Melissa; Dioscoreophyllum* |
| 78508-96-0 | L-erythro-ascorbate | Oxygenated aliphatics | | Not listed in SPECIATE | 10176122 | No | No | Undetermined | Yes | *Taraxacum* |
| 106-70-7 | Methyl hexanoate | Oxygenated aliphatics | | 10 profiles; top sectors: Biomass Burning; Residential Wood Combustion; Outdoor Boiler (6); Biomass Burning; Residential Wood Combustion; Hydronic Heater (3); Agriculture; Silage (1) | 7824 | No | No | Mixed | Yes | *Gmelina; Ginkgo; Syzygium; Calea; Asteriscus; Ligularia; Senegalia; Hypericum (+23 more)* |
| 111-82-0 | Methyl laurate | Oxygenated aliphatics | | 1 profiles; top sectors: Textile Products (1) | 8139 | Yes | Yes | Mixed | Yes | *Helianthus; Vaccinium; Bassia; Crotalaria; Angiopteris; Marchantia; Glycine; Mentha (+15 more)* |
| 6622-76-0 | Methyl tiglate | Oxygenated aliphatics | | Not listed in SPECIATE | 5323652 | No | No | Undetermined | Yes | *Averrhoa; Polyalthia* |
| 96-31-1 | N,n'-dimethylurea | Oxygenated aliphatics (UNRESOLVED) | | Not listed in SPECIATE | 7293 | No | No | Undetermined | No | *nan* |
| 123-39-7 | N-methylformamide | Oxygenated aliphatics (Others (N, S, Halo)) | | Not listed in SPECIATE | 31254 | No | Yes | Anthropogenic | Yes | *Astragalus; Aloe* |
| 925-60-0 | N-propyl acrylate | Oxygenated aliphatics | | Not listed in SPECIATE | 13550 | No | No | Undetermined | – | *nan* |
| 31502-14-4 | Non-2-en-1-ol | Oxygenated aliphatics (UNRESOLVED) | | Not listed in SPECIATE | 5364941 | No | No | Undetermined | Yes | *Gratiola; Centella; Achillea; Gaylussacia; Curcuma* |
| 124-19-6 | Nonanal | Oxygenated aliphatics | | 80 profiles (4 biogenic); top sectors: Mobile; Onroad; Light Duty (26); Mobile; Onroad (24); Agriculture; Animal (5) | 31289 | Yes | Yes | Mixed | Yes | *Syzygium; Strobilanthes; Gmelina; Brunfelsia; Morus; Emblica; Strychnos; Saururus (+71 more)* |

| | | | | | | | | | | |
|---|---|---|---|---|---|---|---|---|---|---|
| 112-05-0 | Nonanoic acid | Oxygenated aliphatics | | 89 profiles (12 biogenic); top sectors: Mobile; Onroad; Light Duty (33); Mobile; Onroad (21); Cooking; Frying (6) | 8158 | Yes | Yes | Mixed | Yes | *Ocotea; Byrsonima; Asparagus; Rudbeckia; Parasenecio; Sonchus; Sambucus; Astragalus (+97 more)* |
| 3391-86-4 | Oct-1-en-3-ol | Oxygenated aliphatics (UNRESOLVED) | | 3 profiles; top sectors: Consumer Products (3) | 18827 | No | No | Mixed | Yes | *Durio; Vaccinium; Opuntia; Ocimum; Solanum; Capsicum; Lathyrus; Koenigia (+80 more)* |
| 2548-87-0 | Oct-2-enal | Oxygenated aliphatics | | Not listed in SPECIATE | 5283324 | No | No | Biogenic | Yes | *Pentanema; Eupatorium; Coriandrum; Ribes; Haplocarpha; Lippia; Cleistopholis; Solidago (+45 more)* |
| 124-13-0 | Octanal | Oxygenated aliphatics | | 88 profiles (4 biogenic); top sectors: Mobile; Onroad (31); Mobile; Onroad; Light Duty (26); Agriculture; Animal (6) | 454 | Yes | Yes | Mixed | Yes | *Brassica; Cucumis; Cymbopogon; Cnidium; Catharanthus; Daucus; Theobroma; Amelanchier (+46 more)* |
| 124-07-2 | Octanoic acid | Oxygenated aliphatics | | 94 profiles (12 biogenic); top sectors: Mobile; Onroad; Light Duty (37); Mobile; Onroad (20); Biomass Burning; Residential Wood Combustion; Woodstove (7) | 379 | Yes | Yes | Mixed | Yes | *Brassica; Cucumis; Cymbopogon; Juglans; Morus; Daucus; Apium; Ribes (+75 more)* |
| 106-36-5 | Propyl propionate | Oxygenated aliphatics | | 4 profiles; top sectors: Agriculture; Silage (4) | 7803 | No | No | Biogenic | Yes | *Fagopyrum; Mentha; Vaccinium; Cocos; Pinus; Galega; Capsicum* |
| 124-25-4 | Tetradecanal | Oxygenated aliphatics | | 2 profiles; top sectors: Cooking; Charbroiling (2) | 31291 | No | No | Biogenic | Yes | *Cebatha; Hedyotis; Capsicum; Morella;* |

| | | | | | | | | | | |
|---|---|---|---|---|---|---|---|---|---|---|
| | | | | | | | | | | *Entada; Yucca; Lupinus; Erucastrum (+49 more)* |
| 637-65-0 | Tetrahydrofurfuryl propionate | Oxygenated aliphatics | | Not listed in SPECIATE | 61183 | No | No | Undetermined | Yes | *Tragopogon* |
| 6138-85-8 | Tetrahydroionone | Oxygenated aliphatics (Terpenoids) | | Not listed in SPECIATE | 110787 | No | No | Undetermined | Yes | *nan* |
| 2277-16-9 | Trans-4-nonenal | Oxygenated aliphatics | | Not listed in SPECIATE | 5283337 | No | No | Undetermined | Yes | *Annona; Mikania* |
| 10486-19-8 | Tridecanal | Oxygenated aliphatics | | 6 profiles; top sectors: Mobile; Onroad (3); Cooking; Charbroiling (2); Mobile; Nonroad (1) | 25311 | No | No | Mixed | Yes | *Koenigia; Capsicum; Artemisia; Zanthoxylum; Lupinus; Conium; Satureja; Musa (+25 more)* |
| 112-44-7 | Undecanal | Oxygenated aliphatics | | 11 profiles (2 biogenic); top sectors: Mobile; Onroad (4); Cooking; Frying (3); Cooking; Charbroiling (2) | 8186 | No | No | Mixed | Yes | *Forsythia; Byrsonima; Asparagus; Vigna; Opuntia; Ananas; Panax; Lathyrus (+56 more)* |
| 112-37-8 | Undecanoic acid | Oxygenated aliphatics | | 70 profiles (7 biogenic); top sectors: Mobile; Onroad (20); Mobile; Onroad; Light Duty (20); Biomass Burning; Residential Wood Combustion; Fireplace (13) | 8180 | No | No | Mixed | Yes | *Vigna; Mentha; Rubus; Fritillaria; Pedicularis; Cocos; Aloe; Beta (+25 more)* |
| 1195-32-0 | P-mentha-1,3,5,8-tetraene | Terpenes | | Not listed in SPECIATE | 62385 | No | No | Biogenic | Yes | *Capsicum; Vanilla; Nelumbo; Origanum; Physalis; Piper; Ribes; Solanum (+58 more)* |
| 127-51-5 | Alpha-isomethylionone | Terpenoids | | Not listed in SPECIATE | 5372174 | No | No | Biogenic | Yes | *Hibiscus; Beta; Grindelia; Senecio; Hedyotis; Daucus; Achillea* |

| | | | | | | | | | | |
|---|---|---|---|---|---|---|---|---|---|---|
| 79507-84-9 | Artemisyl propionate | Terpenoids (Oxygenated aliphatics) | | Not listed in SPECIATE | 101415797 | No | No | Biogenic | Yes | *Allium; Punica* |
| 22339-23-7 | Calamenene | Terpenoids | | Not listed in SPECIATE | 11298625 | No | No | Biogenic | Yes | *Metroxylon; Glycyrrhiza; Houttuynia; Euchresta; Glycine; Andira; Pericopsis; Valerianella (+29 more)* |
| 3796-70-1 | Geranyl acetone | Terpenoids | | 1 profiles; top sectors: Cigarette (1) | 1549778 | No | No | Biogenic | Yes | *Gleditsia; Acacia; Artemisia; Morithamnus; Loxothysanus; Punica; Cicer; Psophocarpus (+51 more)* |

### Supplementary Table S3

Summary of Supplementary Table S2. Number of compounds in each chemical class assigned to each likely-source category for the sampling context of this study. Categories and the evidence used to assign them are defined in the legend to Supplementary Table S2. Counts are of distinct compounds and are not weighted by abundance; terpenes and terpenoids account for 42 of the 262 compounds detected but dominate the vegetation-associated signal reported in the main text.

| Chemical class | Biogenic | Biogenic (dual source) | Mixed | Anthropogenic | Undetermined | Total |
|---|---|---|---|---|---|---|
| Terpenes | 16 | 6 | 0 | 0 | 0 | 22 |
| Terpenoids | 14 | 6 | 0 | 0 | 0 | 20 |
| Oxygenated aliphatics | 12 | 0 | 24 | 3 | 25 | 64 |
| Aromatics | 6 | 0 | 33 | 15 | 18 | 72 |
| Aliphatic hydrocarbons | 2 | 0 | 13 | 19 | 23 | 57 |
| Others | 1 | 0 | 4 | 5 | 17 | 27 |
| **Total** | 51 | 12 | 74 | 42 | 83 | 262 |
